# Evolutionary innovation in floral reproduction by modifications in stem cell peptide signaling

**DOI:** 10.1101/2025.06.27.661788

**Authors:** Daniel S. Jones, Reid Selby, Pedro Jiménez-Sandoval, Andrew C. Willoughby, Krishna Reddy Challa, Emily Yaklich, Anna T. DiBattista, Jakub Baczynski, Feng Wang, Teng Zhang, Vandana Gurung, Ashley D. Crook, Andra-Octavia Roman, Erika R. Moore-Pollard, Riley Schuld, Jennifer R. Mandel, Paula Elomaa, John M. Burke, Julia Santiago, Zachary L. Nimchuk

**Affiliations:** Department of Biology, University of North Carolina at Chapel Hill, Chapel Hill, 27599, U.S.A; Curriculum in Genetics, University of North Carolina at Chapel Hill, Chapel Hill, 27599, USA; Department of Biological Sciences, Clemson University, Clemson, 29634, U.S.A; Department of Biological Sciences, Auburn University, Auburn, 36849, Alabama, U.S.A; The Plant Signaling Mechanisms Laboratory, Department of Plant Molecular Biology, University of Lausanne, Lausanne, 1015, Switzerland; Department of Plant Biology, Miller Plant Sciences, University of Georgia, Athens, 30602, U.S.A; The Plant Center, University of Georgia, Athens, 30602, U.S.A; Institute of Evolutionary Biology, Faculty of Biology, Biological and Chemical Research Centre, University of Warsaw, Warsaw, 02-096 Poland; Department of Agricultural Sciences, Viikki Plant Science Centre, University of Helsinki, Helsinki, 00014, Finland; Department of Biological Sciences, University of Memphis, Memphis, 38152, U.S.A

## Abstract

Understanding how evolution shapes genetic networks to create new developmental forms is a central question in biology. Asteraceae (sunflower family) comprise 10% of flowering plants and have capitula, a novel flowering shoot (inflorescence) that mimics a single flower (*1*, *2*). During capitulum development, shoot stem cells undergo prolonged proliferation relative to other species (*3*, *4*). Here we show that capitulum evolution paralleled decreases in CLAVATA3 peptide (CLV3p) signaling, a conserved repressor of stem cell proliferation. Asteraceae CLV3p displays reduced receptor binding and downstream transcriptional outputs. Reversion of *CLV3* to a more active form impairs Asteraceae stem cell regulation and capitulum development. Lastly, we trace *CLV3* evolution across the Asterales allowing inferences on capitulum evolution. Our findings reveal novel evolutionary mechanisms in plant reproduction and suggest approaches for engineering crops (*5*–*8*).

## Main Text

The deep conservation of kingdom specific developmental genes in animals and plants suggests that evolution acts to fine tune developmental networks rather than replace or eliminate them. The re-wiring of developmental networks through altered cis-regulatory control has been associated with evolutionary novelty across kingdoms, but it is less clear how evolutionary novelty arises when network wiring is conserved. In plants, inflorescence architecture has been highly modified through domestication in diverse crop species and several artificially selected alleles contributing to this have been identified (*9*). In contrast, little is known about the genetic mechanisms that govern major shifts in the natural evolution of inflorescence form across Angiosperms. As the largest family, Asteraceae (sunflower family) account for about 10% of all flowering plants and are defined by a novel inflorescence form called a capitulum, or head (*1*, *10*), as famously depicted in the sunflower paintings of van Gogh. The capitulum is composed of multiple individual florets (flowers) that show considerable diversity in morphology and organization across Asteraceae species (*11*). The aggregated capitulum mimics a solitary flower (pseudanthium) that is thought to contribute to successful pollination and seed dispersal (*10*, *12*), both key traits underlying the global success of Asteraceae. As in other flowering plants, the capitulum is derived from a shoot apical meristem (SAM) that contains a pool of self-maintaining stem cells at the growing apex. In most Angiosperms studied, the size of this pool, and the size of the meristem itself, are relatively constant during inflorescence elaboration, with cells exiting the stem cell zone differentiating into floral primordia or secondary meristems, as in *Arabidopsis thaliana* (*13*, *14*) (Arabidopsis; Fig. 1a). In contrast, capitulum development in Asteraceae is characterized by a prolonged period of stem cell proliferation, and a corresponding increase in meristem size, followed by differentiation of centripetal spirals of florets across the expanded SAM surface, forming a pseudanthium as in *Helianthus annuus* L. (sunflower; Fig. 1b) (*3*, *4*, *15*, *16*). How the Asteraceae capitulum evolved is unclear, but the conservation of this trait across the family reflects its likely monophyletic origin. Closely related families to Asteraceae in the MGCA (Menyanthaceae, Goodeniaceae, Calyceraceae, Asteraceae) clade, exhibit raceme or thyrse-like inflorescence development: flowering shoots with indeterminate meristems, elongated internodes, and lateral flower or branch formation (in Menyanthaceae and Goodeniaceae). In contrast, Calyceraceae (sister to the Asteraceae) have head-like inflorescences called cephalioids, which share the compacted disc-like architecture seen in Asteraceae but maintain terminal flower determinacy and cymose branching patterns of development (*17*). Currently, there are two models describing the evolution of the Asteraceae capitulum. The *inflorescence-origin model* posits that the capitulum evolved from an ancestral thryse-like inflorescence via shoot compaction and SAM expansion, followed by a loss of apical determinacy (loss of a terminal flower) (*17*). In contrast, the *floral-unit-meristem model* proposes that a capitulum meristem is determinate and similar to a floral meristem in structure, as it evolved through paedomorphosis of a floral meristem (*18*). While the origins of the capitulum remain unclear, both models require expanded meristem stem cell proliferation as a critical step in capitulum evolution. Here we show that the evolution of the novel Asteraceae capitulum inflorescence was associated with a quantitative reduction in stem cell repressing peptide signaling, which is critical for normal capitulum development.

**Fig 1:**
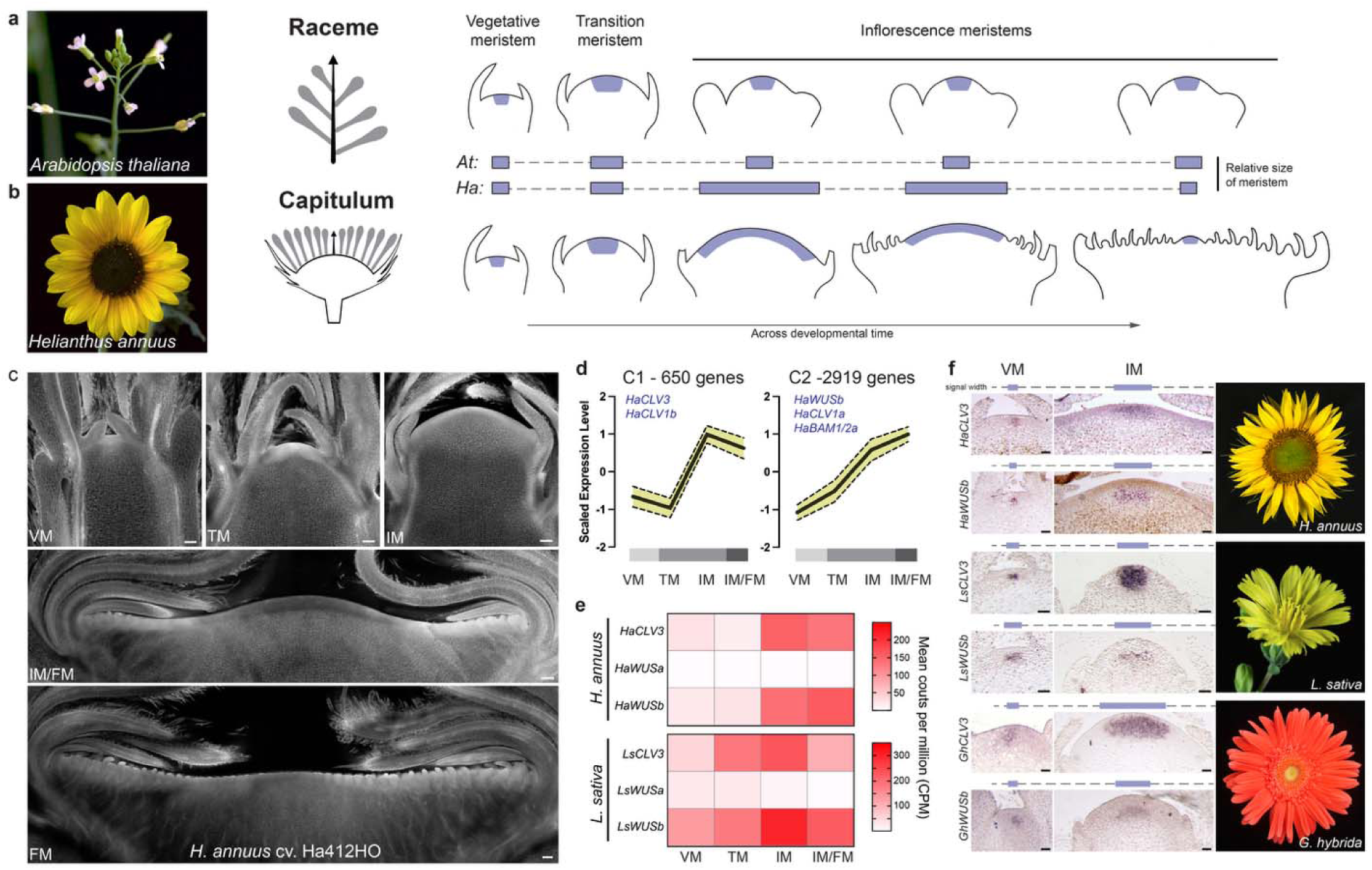
Stem cell proliferation occurs prior to floral initiation in Asteraceae capitula. a-b, Shoot meristem dynamics during inflorescence development in *Arabidopsis thaliana* (raceme) and *Helianthus annuus* (sunflower; capitulum) with relative meristem size in blue. Models based on data from previous studies (*14-16*) c, Confocal micrographs of *H. annuus* cv. Ha412HO shoot meristems spanning floral transition, meristem expansion and floral initiation; representative of stages used for transcriptomics. d, Scaled expression of genes from clusters 1 and 2 following clustering analysis of DE genes (solid line = mean; dotted lines = standard deviation) with total number of genes in each cluster and representative *CLAVATA* pathway genes present indicated. e, Heatmap showing mean expression of *CLV3, WUSa,* and *WUSb* in sunflower and *Lactuca sativa* (lettuce) meristems during inflorescence development. f, *in situ* hybridization *CLV3* and *WUSb* in sunflower, lettuce and *Gerbera hybrida* (Gerbera) at VM and IM stages. Width of *in situ* signal indicated above images with blue line showing expansion of expression domains across species. VM = vegetative meristem, TM = transition meristem, IM = inflorescence meristem, FM = floral meristem. Scale bars - 100µm (c), 50µm (f).

### Asteraceae inflorescences display novel *WUS*-*CLV* stem cell dynamics

To define the genetic requirements of capitulum development, we first used RNA-sequencing (RNAseq) to analyze gene expression in SAMs of cultivated sunflower (*H. annuus* cv. HA412-HO [PI 642777] – reference genome line (*19*)) at key developmental transitions. Stages were identified using confocal microscopy of longitudinal sections from sunflower shoot apices throughout development (Fig. 1c). Three replicates of sunflower SAMs were sequenced spanning vegetative growth (vegetative meristem – VM), at floral transition (transition meristem – TM), during meristem expansion (inflorescence meristem – IM), and early floret initiation (inflorescence and floral meristems – IM/FM). Differential expression analysis identified 6,657 (VM x TM), 8,265 (TM x IM), and 2,095 (IM x IM/FM) significantly differentially expressed genes (DEGs) during capitulum development (Supplementary Fig. 1a-b; Supplementary File 1). We used a clustering analysis to investigate generalized patterns of gene co-expression across the developmental series, using the 12,657 DEGs identified from pairwise comparisons (Supplementary Fig. 1c-d; Supplementary File 1). Thirteen clusters were identified using a hierarchical clustering-derived cutoff to define groups with similar expression patterns (Supplementary Fig. 1c). Clusters 1 (650 genes) and 2 (2,919 genes) exhibited increased gene expression during capitulum meristem expansion and early floral initiation phases (Fig. 1d). Consistent with these developmental stages, both clusters contained orthologs of key meristem regulatory genes including members of the highly conserved *CLAVATA* (*CLV*) signaling pathway (Supplementary File 1).

In Angiosperms, shoot stem cell proliferation is repressed by CLAVATA3 peptide signaling (CLV3p), the founding member of the CLAVATA3/EMBRYO-SURROUNDING REGION peptide (CLEp) hormone family (*20*–*22*). Mutations in *CLV3* lead to uncontrolled stem cell proliferation, enlarged shoot size (fasciation), and increases in floral organ numbers. The mature processed 12 amino acid CLV3p is derived from the CLE domain of the CLV3 apoprotein, secreted from shoot stem cells, and diffuses into the meristem center where it signals through the CLAVATA1 (CLV1) leucine rich repeat (LRR) receptor kinase, and accessory receptors, to repress expression of the stem cell promoting transcription factor *WUSCHEL* (*WUS*) (*13*, *23*). In Arabidopsis, loss of *CLV1* is partially compensated by transcriptional upregulation of the CLV1-related *BARELY ANY MERISTEM 1/2* (*BAM1/2*) receptors, which then participate in CLV3p-dependent stem cell regulation in all shoot meristem types (*21*, *24*, *25*). We leveraged recently published genomic resources and generated new *de novo* transcriptome assemblies of SAMs in six species from Asteraceae and immediate outgroup taxa to identify *CLV* pathway orthologs across the Asterales (Supplementary Fig. 2) (*26*). Ortholog analyses consisted of data from major Angiosperm clades (ANA: 2 species, Magnoliid: 3 sp., Monocots: 12 sp., and Eudicots: 39 sp.), including a diverse set of Asteraceae spanning 5 subfamilies and 13 tribes (Supplementary File 2; Supplementary Fig. 2). Through these analyses, we identified Asteraceae-specific duplications of *CLV1*, *BAM1/2*, and *WUS* (Supplementary Fig. 3). We then used two previously reported Hidden Markov Model (HMM) profiles of CLE motifs to identify and classify Asteraceae *CLEs* (Supplementary File 3) (*27*, *28*). Clustering analysis based on sequence similarity revealed only one CLE1D member (from the 1D1 subclade) in all Asteraceae species examined (16 total), indicating that a single *CLV3* ortholog is present in all Asters. This ortholog groups with genetically defined *CLV3* genes from Arabidopsis and tomato (Fig. 2a; Supplementary Fig. 4) (*27*, *29*). We further validated both *CLV3* and *CLV1* ortholog identification using a directed BLAST (tblastn) to compare sunflower and *Lactuca sativa* L. (lettuce; Asteraceae) orthologs against eight genomes (four Asteraceae and four from closely related outgroups) (*26*). We found that both sunflower and lettuce CLV3 return only one high similarity hit in all genomes surveyed and this analysis also supported the existence of the Asteraceae-specific *CLV1* duplication (Supplementary Files 4-5).

**Fig 2:**
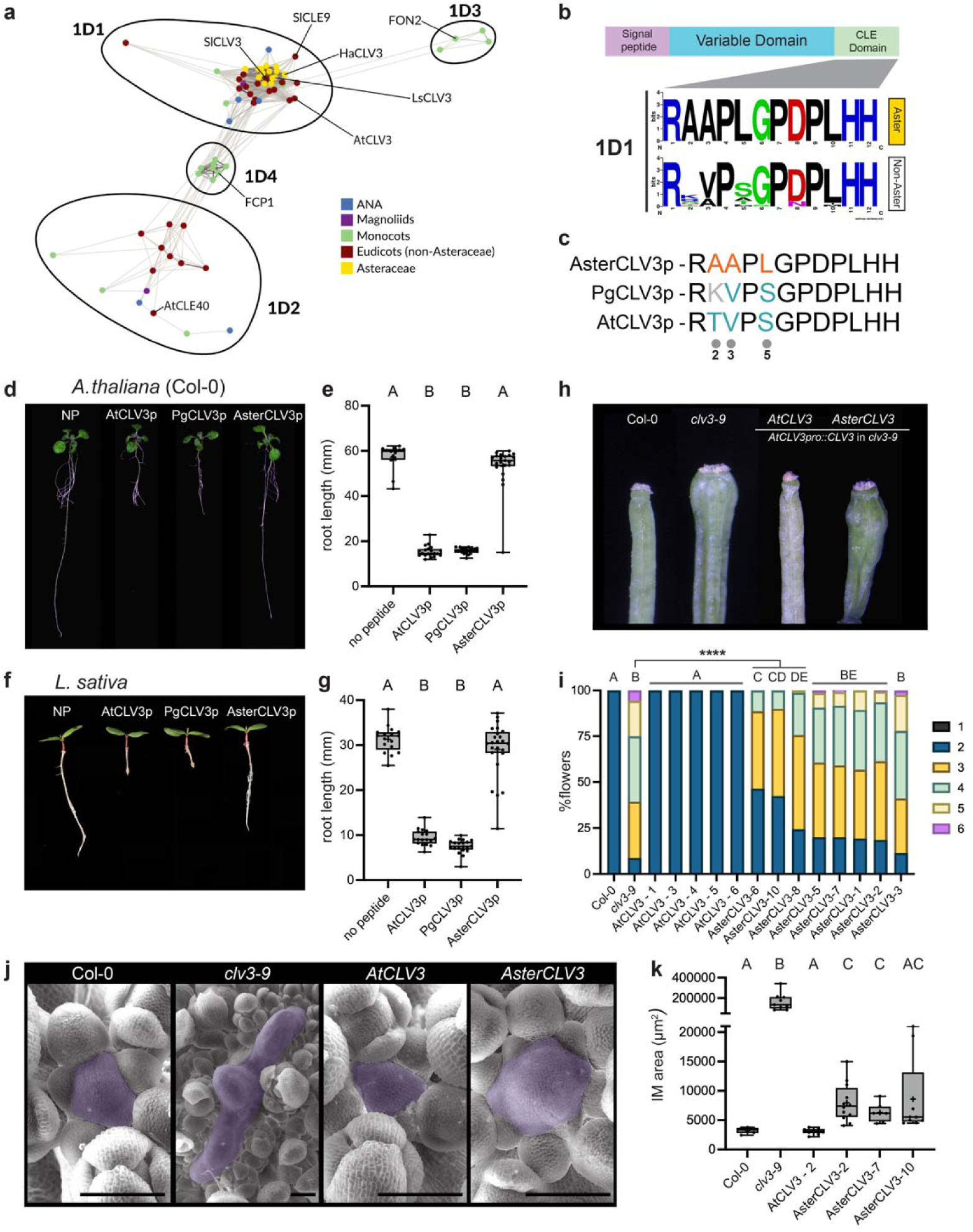
Reduced signaling activity of Asteraceae *CLV3*. a, Clans-based clustering of CLE1D-type *CLEs* from across flowering plant lineages with genes from *A. thaliana* (At), *Solanum lycopersicum* (Sl), *Oryza sativa* (*FCP1* and *FON2*), sunflower and lettuce labeled. b, LOGO consensus diagrams of mature CLE domains from all CLE1D1-type CLEs showing conservation of CLE motif in all Asteraceae *CLV3* orthologs analyzed. c, CLV3p CLE domain sequence from Asteraceae (Aster), *Platycodon grandiflorus* (Pg; from Campanulaceae, an Asteraceae outgroup) and Arabidopsis (At) with residues 2, 3 and 5 labeled at bottom. d-g, CLE peptide root elongation assays showing responses to 0.1μM AtCLV3p, PgCLV3p and AsterCLV3p in *A. thaliana* (d-e) and lettuce (f-g) each compared to no peptide (NP) controls. Statistical summary from one-way ANOVA where A and B groups are significantly different (adj. p-value <0.0001). h-i, T3 complementation analysis of carpel number in *clv3-9* using a native *AtCLV3* promoter::gene (*AtCLV3pro::AtCLV3*) constructs with AtCLV3 and AsterCLV3 CLE domains (related to Supplementary Fig. 6d). Statistical summary from Kruskal-Wallis and Dunn’s multiple comparison tests where A-E groups are significantly different (adj. p-value <0.05) with three AsterCLV3 lines (6, 10, and 8) highlighted as significantly different than *clv3-9* (**** = adj. p-value <0.0001) (related to Supplementary Fig. 6d). j-k, Scanning electron micrographs (SEM) of IMs from representative Arabidopsis complementation lines (*AtCLV3pro::AtCLV3* and *AtCLV3pro::AsterCLV3* in *clv3-9* background) and controls along with quantification of IM area (µm^2^). Statistical summary from Brown-Forsythe and Welch ANOVA where A-C groups are significantly different (adj. p-value <0.05). Varying residues labeled in orange (if similar to AsterCLV3p) and blue (if similar to AtCLV3p).In e, (n=individual roots analyzed) n= 13 (NP), 20 (AtCLV3p), 21 (PgCLV3p) and 21 (AsterCLV3p). In g, (n=individual roots analyzed) n= 17 (NP), 18 (AtCLV3p), 20 (PgCLV3p) and 24 (AsterCLV3p). In i, (n=individual plants with 18-20 flowers counted on each) n= 6-8 plants. In k, (n=individual IMs measured) n= 6 (Col-0), 9 (*clv3-9*), 9 (*AtCLV3-2*), 12 *AsterCLV3-2*), 7 (*AsterCLV3-7*) and 9 (*AsterCLV3-10*). See complete statistical analyses and sample sizes in Supplementary File 8. Scale bars - 100µm (j).

In all angiosperms studied to date, elevated *CLV3* expression represses *WUS,* and thereby stem cell proliferation, in a *CLV1*-dependent manner (*13*, *21*, *23*). In contrast, sunflower orthologs of *CLV3* (*HaCLV3*)*, CLV1* (*HaCLV1a/b*)*, WUS* (*HaWUSb*), and *BAM1/2* (*HaBAM1/2a*) were all present in SAM expression clusters 1 and 2, displaying coincident and increased expression during capitulum meristem expansion (Fig. 1d). We next explored the conservation of *CLAVATA* pathway regulation during capitulum meristem expansion using a publicly available RNAseq dataset from similar stages of lettuce SAMs (Fig. 1e) (*30*). Again, *CLV3* and *WUS* (*WUSb*) expression were coincident and peaked in inflorescence meristems during the expansion phase in lettuce, while *WUSa* had low or no detectable expression at the meristem stages sampled (Fig. 1e). In Arabidopsis and tomato, *CLV3* is expressed in central zone stem cells, with *WUS* expression being more prominent in the subtending organizing center (*13*, *23*). In Arabidopsis, *CLV1* is expressed in the meristem center, with lower levels in the *WUS*-expressing organizing cells (*31*), and low expression levels in L1 cells (*32*, *33*). To further confirm Asteraceae *CLV3* orthology we examined the expression domains of *CLV3*, *CLV1a/b* and *WUSb* genes in select Asteraceae models using *in situ* hybridization across meristem stages and types. As in other species, Asteraceae *CLV3* is expressed in the central zone regions of VMs and IMs in representative species from diverged Asteraceae subfamilies; sunflower (*HaCLV3,* Asteroideae), lettuce (*LsCLV3,* Cichorioideae), and *Gerbera hybrida* (Gerbera, *GhCLV3,* Mutisioideae), and *WUSb* is expressed in the organizing center in all three species (Fig. 1f and Supplementary Fig. 5a) (*4*). We further verified this spatial patterning of *CLV3* in a developing capitulum using a native *GhCLV3pro* reporter (*GhCLV3pro::NLS-eGFP*) line in Gerbera (Supplementary Fig. 5b). *CLV1a/b* were expressed in both VMs and IMs across all species as well, with expression being highest in apical regions of the meristems and decreasing towards the organizing center (Supplementary Fig. 5c-e). Notably, the size of the expression domains of *CLV3*, *WUSb*, and *CLV1a/b* increased in all three species during the VM to IM transition, paralleling the expansion of the capitulum. (Fig. 1f; Supplemental Fig. 5). As such, while there is conservation of the expression patterns of the core *CLV3*-*WUS* genes in Asteraceae, continuous expression of these genes are not associated with buffered SAM size as in other plant species, suggesting a possible reduction in CLV-signaling in Asteraceae.

### Asteraceae evolved a unique CLV3 peptide variant with reduced signaling capacity

The duration of stem cell proliferation during capitulum development is prolonged relative to other species but not unrestricted as in *clv* mutants, suggesting a quantitative change in stem cell regulation occurred during Asteraceae evolution. Given the importance of CLV3p in stem cell repression, and the apparent differences in *CLAVATA* expression and regulatory dynamics in Asteraceae, we investigated whether capitulum development and evolution could be associated with reductions in CLV3p signaling. To explore if this is due to differences in CLV3p itself, we first compared the CLE motif of Asteraceae and non-Asteraceae *CLV3* homologs (CLE1D1-type *CLEs*), as this represents the mature CLV3p (Fig. 2b). Surprisingly, this revealed that CLV3p is invariant across all 16 Asteraceae species examined. Additionally, the N-terminus of AsterCLV3p diverged significantly from non-Asteraceae CLV3p, for which position 2 is variable and there is a bias towards valine and serine at positions 3 and 5, respectively (for CLE1D1s). In contrast these residues in AsterCLV3p were invariably Ala2, Ala3, and Leu5, forming an N-terminal CLEp motif we term the RAAPL CLV3p (Fig. 2b). To test if the AsterCLV3p displayed altered activity, we first performed root growth inhibition assays, a standard measure for CLEp activity (*34*), in response to CLV3p peptides from Arabidopsis AtCLV3p (RTVPS), AsterCLV3p (RAAPL), and from *Platycodon grandiflorus* PgCLV3p (RKVPS, Campanulaceae), an outgroup to the MGCA clade (Fig. 2c). While AtCLV3p and PgCLV3p repressed root growth in Arabidopsis seedlings at low peptide concentrations, AsterCLV3p failed to do so (Fig. 2d-e). This was mirrored in lettuce and sunflower seedling root assays, revealing that Asteraceae can respond to CLV3p from other species but fail to respond to their own in this assay (Fig. 2f-g; Supplementary Fig. 6a). We next tested the ability of AsterCLV3p to promote *CLV1*-dependent stem cell repression in shoots, first using Arabidopsis as a proxy model. CLV3-CLV1 signaling operates in FMs and IMs to maintain floral organ number and meristem size respectively. Consistent with the root growth inhibition assays, expression of either *AtCLV3* (RTVPS) or full-length *AtCLV3* containing *PgCLV3* CLEp motif substitutions (RKVPS), under the native *AtCLV3* promoter (*AtCLV3pro::CLV3var*), fully complemented Arabidopsis *clv3* (*clv3-9*) carpel number increases while the *AsterCLV3* CLEp motif (RAAPL) substitutions provided an intermediate level of complementation, with 3/8 T3 homozygous lines being significantly different from *clv3-9* (Fig. 2h-i, Supplementary Fig. 6d-e). While multiple independent homozygous single transgene insertion lines displayed significant reductions in carpel numbers compared to *clv3*, this effect was not significant in pooled analysis of *AsterCLV3* expressing T1 lines (Fig. 2h-i). Analysis of total *CLV3* transcript levels in lines with confirmed *AtCLV3pro::CLV3var* transgene expression revealed an intermediate impact on the feedback-repression of *CLV3* in *AsterCLV3* lines compared to *AtCLV3* lines, consistent with quantitatively reduced AsterCLV3p function (Supplementary Fig. 6f-g). This intermediate activity for *AsterCLV3* was also mirrored in IMs, with *AtCLV3* restoring wild-type IM size to *clv3* mutants, as expected from previous data (*35*), and multiple homozygous *AsterCLV3* lines providing an intermediate degree of complementation (Fig 2j-k). Collectively these data indicate that AsterCLV3p has reduced *CLV1*-dependent stem cell–restricting activity in both Arabidopsis FMs and IMs (Fig. 2h-k; Supplementary Fig. 6d).

In Arabidopsis, CLV1 and its close homologs BAM1/2/3 mediate developmental responses to diverse CLE peptides beyond the shoot apical meristem, whereas PXY (PHLOEM INTERCALATED WITH XYLEM) receptors specifically recognize TDIF (TRACHEARY ELEMENT DIFFERENTIATION INHIBITORY FACTOR) class CLE peptides (CLE41p, CLE42p, CLE44p) to regulate vascular development (*36*, *37*). CLE peptide signaling requires co-receptors (*38*–*40*); however, direct and biologically relevant CLEp binding has been only shown for CLV1, BAM, and PXY receptors, with specificity and affinity varying across receptor–ligand pairs (*41*–*46*). Although all belong to the same LRR-RLK subfamily, structural predictions and phylogenetic analyses reveal distinct architectures of the ligand-binding pocket between CLV1/BAM and PXY receptors, consistent with their separation into different evolutionary subclades (Supplementary Fig. 7) (*47*). Despite this, genetic, biochemical, and structural studies—based on the PXY–TDIF complex—have identified a conserved mechanism of CLE peptide perception across receptor and peptide subclades (*43*, *46*). CLE peptide recognition is primarily mediated by the N-terminal region, which docks the peptide into the receptor pocket through hydrophobic and electrostatic interactions, while the conserved central and C-terminal residues stabilize the complex by facilitating a conformation compatible with the receptor binding groove (*43*, *46*). Although no structures of CLV1/CLV3p complexes have been reported in any species, structural modeling and functional analysis supports a similar binding mode for AtCLV3p and AtCLV1, with essential N-terminal (Arg1, Pro4, Gly6) and C-terminal residues (Asp8, Pro9, His11, His12) being required for *AtCLV3* function *in vivo* (*48*). In AsterCLV3p, conservation of these core N- and C-terminal residues indicates the potential for interaction with CLV1-class receptors (Fig. 2b). Consistent with this, higher doses of AsterCLV3p modestly inhibited Arabidopsis root elongation, suggesting that AsterCLV3 has strongly attenuated, but not eliminated, activity (Supplementary Fig. 6b-c). AsterCLV3p differs from AtCLV3p, and other CLV3p orthologs, at N-terminal positions 2, 3, and 5 (Ala2, Ala3, and Leu5 in AsterCLV3p). Alanine substitution experiments of these positions individually in *AtCLV3* previously suggested they play negligible roles in *CLV1*-dependent stem cell regulation (*48*). This implies that the reduced activity of AsterCLV3p arises from the unique combination of these substitutions. Indeed, the reduction in AtCLV3p activity in sunflower, lettuce and Arabidopsis root length peptide sensitivity assays was strongest when Ala2, Ala3, and Leu5 substitutions were combined (Fig. 3a-b, Supplementary Fig. 8a). Moreover, these substitutions were additive in attenuating *CLV1*-dependent stem cell repression in Arabidopsis shoots across T1 plants, as measured by carpel numbers, with *AtCLV3^Ser5Leu^* variants having a negligible effect on activity on their own but additive effects with the *AtCLV3^Thr2Ala, Val3Ala^* substitutions (thereby recreating the AsterCLV3p RAAPL sequence; Fig. 3c, Supplementary Fig. 8b-c). We observed similar patterns of partial complementation of *clv3-9* for increased rosette leaf number and stem fasciation phenotypes, with the *CLV3* variants tested having intermediate effects, including *AsterCLV3* (Supplementary Fig. 8d-f). As such, the strongly attenuated activity of AsterCLV3p is due to a unique combination of substitutions outside of the essential conserved CLE peptide residues and suggest that Ser5 is an important residue for CLV3p recognition. As Arabidopsis BAM1/2 participate with CLV1 in CLV3p-dependent shoot stem cell regulation we next asked if AsterCLV3p substitutions impact BAM1/2 signaling specifically. To do this we assayed the ability of CLE peptide variants to promote formative divisions in root ground tissue layers (cortex and endodermal layer), as this requires *BAM1/2* but is independent of *CLV1* (*49*). The number of ground tissue layers were scored using the *EN7* reporter, which marks the inner endodermal layer (*50*). Both AtCLV3p and AtCLE13p, a relevant root BAM1 CLE-ligand (*49*), promoted division in root ground tissues giving rise to additional ground layers beyond the invariant two in untreated roots (Fig. 3d). In contrast, AsterCLV3p had no effect on ground tissue divisions compared to no peptide controls. Interestingly, Ser5Leu substitutions in both AtCLV3p and AtCLE13p diminished activity, while Leu5Ser substitutions enabled AsterCLV3p to induce *BAM1/2*-mediated ground tissue divisions. Leu5 substitutions also impaired the activity of 8 out of 9 tested Arabidopsis CLE peptides in root growth inhibition assays (Fig. 3e). Collectively these data support a general requirement for position 5 in effective CLE peptide– receptor interactions.

**Fig 3:**
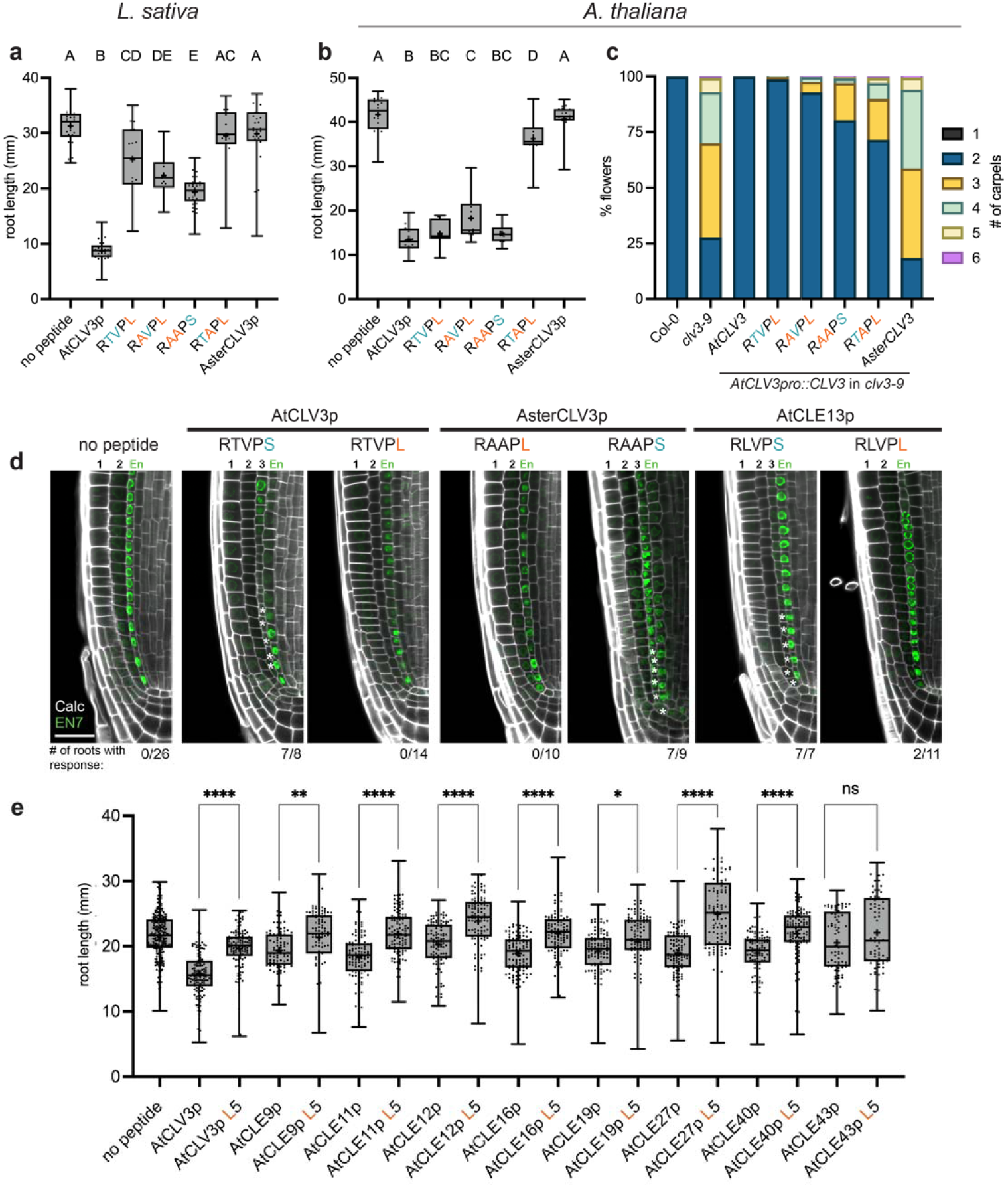
Contributions of N-terminal residues to AsterCLV3 signaling. a-b, CLE peptide root elongation assays showing responses to 0.1μM AtCLV3p, AsterCLV3p and intermediate CLV3p variants in lettuce (a) and *A. thaliana* (b) each compared to no peptide (NP) controls. Statistical summary from one-way ANOVA where A-E groups are significantly different (adj. p-value <0.01). c, T1 complementation analysis of carpel number in *clv3-9* using a native *AtCLV3* promoter::gene (*AtCLV3pro::AtCLV3*) constructs with AtCLV3, RTVPL, RAVPL, RAAPS, RTAPL and AsterCLV3 CLE domains swapped in (related to Supplementary Fig. 8c). d, Root meristem micrographs (*A. thaliana*) of an endodermal marker (green; EN7) with no peptide, or grown in the presence of AtCLV3p, AtCLV3p S5L, AsterCLV3p, AsterCLV3p L5S, AtCLE13p, AtCLE13p S5L. Number of cell layers from the endodermis to the epidermis numbered above and the number of roots with ground tissue responses indicated below micrographs. e, CLE peptide root elongation assay in Arabidopsis showing responses to 0.1μM AtCLEp (with 5^th^ residue Ser) and each AtCLEp S5L variant. Statistics from one-way ANOVA with significance indicated by asterisks. (adj. p-values: *= <0.05, ** = <0.005,**** = <0.0001). Varying residues labeled in orange (if similar to AsterCLV3p) and blue (if similar to AtCLV3p). a-b (n=individual roots analyzed), in a, n= 30 (NP), 31 (AtCLV3p), 20 (RTVPL), 18 (RAVPL), 38 (RAAPS), 13 (RTAPL), and 32 (AsterCLV3p). In b, n= 21 (NP), 21 (AtCLV3p), 10 (RTVPL), 10 (RAVPL), 10 (RAAPS), 11 (RTAPL), and 20 (AsterCLV3p). In c, (n=individual plants with 15 flowers counted each) n= 20 (Col-0), n= 20 (*clv3-9*), 20 (*AtCLV3*), 33 (*RTVPL*), 30 (*RAVPL*), 25 (*RAAPS*), 21 (*RTAPL*), and 26 (*AsterCLV3*). In e (n=individual roots analyzed), n= 252 (NP), 127 and 115 (AtCLV3p and AtCLV3p S5L), 88 and 91 (AtCLE9p and AtCLE9p S5L), 118 and 120 (AtCLE11p and AtCLE11p S5L), 121 and 122 (AtCLE12p and AtCLE12p S5L), 118 and 116 (AtCLE16p and AtCLE16p S5L), 117 and 124 (AtCLE19p and AtCLE19p S5L), 122 and 119 (AtCLE27p and AtCLE27p S5L), 116 and 114 (AtCLE40p and AtCLE40p S5L), 76 and 75 (AtCLE43p and AtCLE43p S5L). See complete statistical analyses and sample sizes in Supplementary File 8. Scale bar - 25μm (d).

We next asked if there were potential changes in the CLEp binding pocket of Asteraceae CLV1a/b orthologs paralleling the altered AsterCLV3 peptide. The N-terminal hydrophobic CLEp binding pocket in CLV1/BAM/PXY receptors is shaped in part by a conserved glycine in the 6th leucine rich repeat (LRR6) which accommodates Val3 in CLE peptides (Supplementary Fig. 9a-d) (*46*). This glycine is essential across CLEp receptor classes and is mutated in the strong EMS-derived *clv1-4* allele in Arabidopsis (AtCLV1^G201E^) (*51*). CLEp substitutions at position 3 with bulky residues would create steric clashes with the receptor LRR6 glycine and other neighboring residues, while CLEp *Val3Ala* substitutions, as in AsterCLV3p, are tolerated (Supplementary Fig. 9) (*42*, *46*, *52*). Inspection of CLV1a orthologs in Asteraceae revealed that this critical LRR6 glycine is substituted by alanine in the majority of species analyzed and is glutamine in lettuce LsCLV1a (LsCLV1a^Q209^) and other species from its tribe (5 species in the Cichorieae; Supplementary Fig. 10a). For CLV1b orthologs, only 4 out of 34 Asteraceae examined had CLV1b LRR6 glycine substitutions, and those were all from species that had additional *CLV1b* paralogs which retain the LRR6 glycine. All CLV1a/b orthologs retain other conserved critical residues implicated in CLEp binding (Supplementary Fig. 10a). Only three Asteraceae species analyzed possessed CLV1a/b orthologs where both retained the canonical LRR6 glycine: sunflower (Heliantheae), *Smallanthus sonchifolius* (Millerieae), and *Mikania micrantha* (Eupatorieae); all closely related tribes from the Heliantheae alliance. Structure prediction with AlphaFold 3 (*53*) revealed that replacement of glycine with alanine at LRR6 reduces the volume of the hydrophobic peptide-binding pocket, potentially weakening binding, while glutamine substitution would introduce steric clashes that are predicted to block CLEp binding, consistent with the defects observed in *Atclv1-4* mutants (Supplemental Fig. 9). Supporting these predictions, native promoter expression of *AtCLV1^G201A^* (*AtCLV1pro::CLV ^G201A^*) and *AtCLV1^G201Q^*substitutions failed to complement Arabidopsis *clv1-101* null mutants, with *AtCLV1^G201A^* lines displaying milder phenotypes than *AtCLV1^G201Q^* lines (Supplementary Fig. 10b-d, Supplementary Fig. 9). Additionally, both *AtCLV1^G201A^* and *AtCLV1^G201Q^*variably enhanced carpel numbers in the *clv1-101* background, mirroring the semi-dominant nature of the *clv1-4^G201E^* allele (Supplementary Fig. 10c) (*20*). To test the conservation of this effect, we generated the same amino acid swaps in the LRR6 glycine of the related AtBAM1 receptor. Native promoter *AtBAM1^G199Q^* (*AtBAM1pro::AtBAM1^G199Q^*) failed to complement the *bam1/2* double mutant male sterility and rosette phenotypes (*54*) (7/7 lines examined), while *AtBAM1* and *AtBAM1^G199A^* complemented both phenotypes (5/5 and 4/4 lines examined, respectively), suggesting AtBAM1^G199A^ retains affinity for CLE peptides (Supplementary Fig. 10e-f). As such, along with the evolution of attenuated AsterCLV3p activity, diverse Asteraceae evolved *CLV1* paralogs with apparent reductions in CLV3p binding potential, while still retaining at least one copy of *CLV1* with predicted CLV3p binding capabilities and signaling potential.

### A novel combination of substitutions in Asteraceae CLV3 peptide reduce receptor affinity

We next investigated whether the reduced activity of AsterCLV3p reflected altered ligand affinity for CLV1/BAM receptors *in vitro*. Despite several attempts and diverse methods, we were unable to purify enough folded AtCLV1 or LsCLV1b proteins. We therefore used AtBAM1 as a proxy receptor for CLEp binding in Isothermal Titration Calorimetry (ITC) assays, due to the conservation of its CLEp-binding pocket with CLV1-type receptors, established CLEp partners, responsiveness to AtCLV3p *in vivo*, role in CLV3p-mediated shoot stem cell regulation (*21*, *24*), and validated AtCLV3p binding *in vitro* (Fig. 3d, Fig. 4a-b, Supplemental Fig. 11) (*43*, *46*, *49*). To determine whether this binding pattern extends to other BAM receptors, we performed ITC assays with AtBAM2 and BAM1/2a from lettuce (LsBAM1/2a). ITC assays were performed to compare binding of the three BAM receptor ectodomains to AtCLV3p, AsterCLV3p, and the high-affinity AtBAM1 ligand AtCLE13p (*49*). The latter was again selected based on its ability to promote BAM1/2-dependent formative cell divisions in Arabidopsis roots (Fig. 3d) (*49*), providing a biologically relevant readout for binding activity. Although AtCLV3p bound AtBAM1 and AtBAM2 with ∼800-fold and ∼10-fold lower affinity than AtCLE13p respectively, both CLE peptides retained robust activity in promoting BAM1/2- dependent Arabidopsis root cell divisions (Fig. 3d). In contrast, AsterCLV3p showed no detectable binding to either AtBAM1 or AtBAM2, mirroring the inability to induce root ground tissue cell divisions (Fig. 3d; Fig. 4c; Supplementary Figs. 12 and 13). Among the residues that diverge from AtCLV3p, position 5 has not previously been implicated in receptor binding or activity. In our refined AtBAM1-CLE13p structural model, the hydroxyl group of CLE13p Ser5 is oriented towards the receptor interface, where it may engage in water-mediated interactions that stabilize the peptide-receptor complex (Fig. 4d-e, Supplementary Fig. 9e; Supplementary Fig. 11f). Supporting this model, substitution of CLE13p Ser5 with Ala or Leu—as found in AsterCLV3p—markedly reduced AtCLE13p binding affinity to AtBAM1/2 (Fig. 4c; Supplementary Figs. 12a-b and 13a), and CLE13p^Ser5Leu^ and AtCLV3p^Ser5Leu^ displayed diminished ability to promote *BAM1/2*-dependent root cell divisions (Fig. 3d). Notably, in the PXY/TDR-CLE41p complex, the equivalent Ser5 side-chain is solvent-exposed and does not contribute to the receptor-peptide binding, suggesting a clade-specific role (*46*). Replacing Leu5 with Ser in AsterCLV3p restored partial binding to AtBAM1/2, but to a level considerably lower than AtCLV3p or AtCLE13p, again suggesting that additional residues—particularly at positions 2 and 3—contribute to full receptor engagement (Fig. 4c. and Supplementary Figs. 12a-b and 13c). Consistent with this AsterCLV3p^Leu5Ser^ displayed activity across bioassays, with significant root growth inhibition activity in sunflower, lettuce, and Arabidopsis (Fig. 3a-b and Supplementary Fig. 8a), a partial ability to complement *clv3* carpel numbers in Arabidopsis (Fig. 3c), and the ability to promote BAM1/2-dependent root cell divisions (Fig. 3d). LsBAM1/2a CLEp binding capabilities mirrored those of AtBAM receptors, with higher affinity binding to AtCLE13p than AtCLV3p, no detectable binding to AsterCLV3p, and with Ser5 playing an important role in promoting binding (Fig. 4c and Supplementary Figs. 12b and 13). Together, these results identify position 5 as an important residue that modulates CLEp activity and receptor binding, contrasting with the role of TDIF Ser5 in PXY receptor interactions. Consistent with our structural model, we observed no binding of any peptide to AtBAM1^G199Q^, while AtBAM1^G199A^ exhibited reduced binding affinities for most CLE peptides and variants tested, mirroring the predicted role of CLEp position 3 in BAM1 interactions (Supplementary Figs. 9 and 12). Notably, neither AtBAM1^G199A^ nor AtBAM1^G199Q^ restored binding to AsterCLV3p (RAAPL), indicating that LRR6 glycine substitutions in Asteraceae species cannot compensate for AsterCLV3p-specific changes (Fig. 4a and Supplementary Figs 9 and 12a-b). Collectively, these findings indicate that the unique and conserved substitutions in AsterCLV3p impair its signaling capacity by reducing receptor binding affinity.

**Fig. 4.**
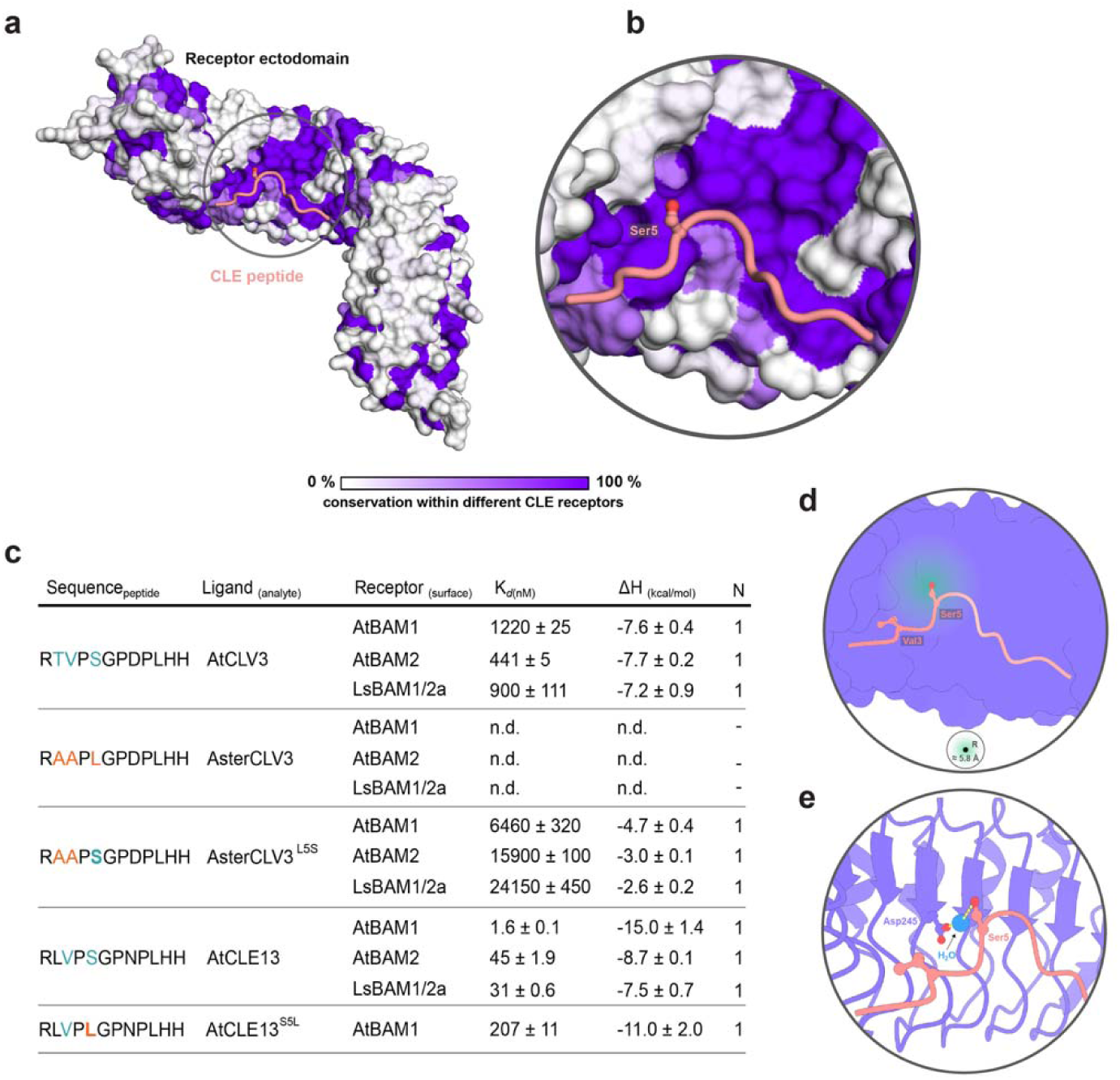
The N-terminal region in CLE peptides impacts BAM/CLV receptor recognition and specificity. a, Surface representation of BAM1/CLV receptor ectodomains, colored based on sequence conservation across *A. thaliana*, sunflower and lettuce (related to Supplementary Fig. 11e). b, Magnified view of the AtBAM1 binding pocket (surface representation colored according to conservation) in complex with CLE13p (pink). c, ITC summary table of CLE/CLV peptides binding to AtBAM1, AtBAM2, and LsBAM1/2a ectodomains. K*_d_*(nM) represents binding affinity, ΔH (kcal/mol) indicates reaction enthalpy, and N denotes stoichiometry (n = 1 for a 1:1 interaction). Data are mean ± SD from at least two independent experiments, with undetected binding shown as n.d. (related to Supplementary Figs. 12 and 13) d-e, CLE13p residues at positions 3 and 5 fit into subpockets on the receptor’s surface, with the Ser5 side chain positioned towards the receptor. Due to the proximity (5.8Å), Ser5 may facilitate water-mediated interactions between the peptide and receptor. The local environment surrounding CLE13p Ser5 supports a network of potential water-coordinated interactions. Potential interactions are indicated by yellow dashed lines (related to Supplementary Fig. 11f). e, Close-up view of a computationally predicted water molecule bridging interactions between CLE13p Ser5 and the receptor binding pocket. The models are based on the AlphaFold3 prediction of AtBAM1 in complex with AtCLE13p.

### Capitulum development and stem cell regulation in Asteraceae requires dampened CLV3p signaling

While we were unable to detect AsterCLV3p binding in our ITC assays, several observations suggest AsterCLV3p retains low levels of signaling capacity *in vivo*. First, AsterCLV3p can restrict Arabidopsis root growth at micromolar concentrations, but not at nanomolar concentrations like AtCLV3p or PgCLV3p (Supplementary Fig. 6b-c). Second, *AsterCLV3* partially suppressed *clv3* carpel numbers in multiple homozygous T3 lines (Fig. 2h-i), and partially suppressed *clv3* IM size, increased leaf numbers, and stem fasciation (Fig. 2j-k; Supplementary Fig. 8d-f). Third, stem cell proliferation is prolonged during capitulum development, but not unrestricted as in *clv3* mutants in other Angiosperms (*21*). We therefore hypothesized that Asteraceae may have evolved a CLV3p with attenuated but not eliminated signaling capacity that permits prolonged stem cell division during capitulum development but still buffers against run-away stem cell proliferation. If so, this would also explain the invariance of the AsterCLV3p across the family and the retention of essential CLV1/BAM binding residues. To test this and determine the function of AsterCLV3 *in vivo*, we generated CRISPR induced *LsCLV3* knockout mutants in lettuce (*lsclv3-cr*) and *GhCLV3* RNAi knockdown lines in Gerbera (Supplementary Fig. 14a-d). *lsclv3-cr* mutants displayed enhanced capitulum and capitulescence (capitula bearing shoot axis) meristem size, stem fasciation and increased floral organ numbers, which we quantified in both petals and stigmata (a proxy for carpel number; Fig. 5a-c and Supplementary Fig. 14e-j). *lsclv3-cr* also developed clusters of fused radial florets in the capitulum center, which we called terminal florets, a phenotype of unknown origin not seen in *clv* mutants in other species (Fig. 5b; Supplementary Fig. 14g). Similar phenotypes were seen in Gerbera *GhCLV3* RNAi knockdown lines, disrupting inflorescence architecture along with increases in meristem size and floral organ numbers (Fig. 5d-e). Together, these data demonstrate that the canonical role for AsterCLV3p in repressing stem cell proliferation is conserved in Asteraceae.

**Fig. 5:**
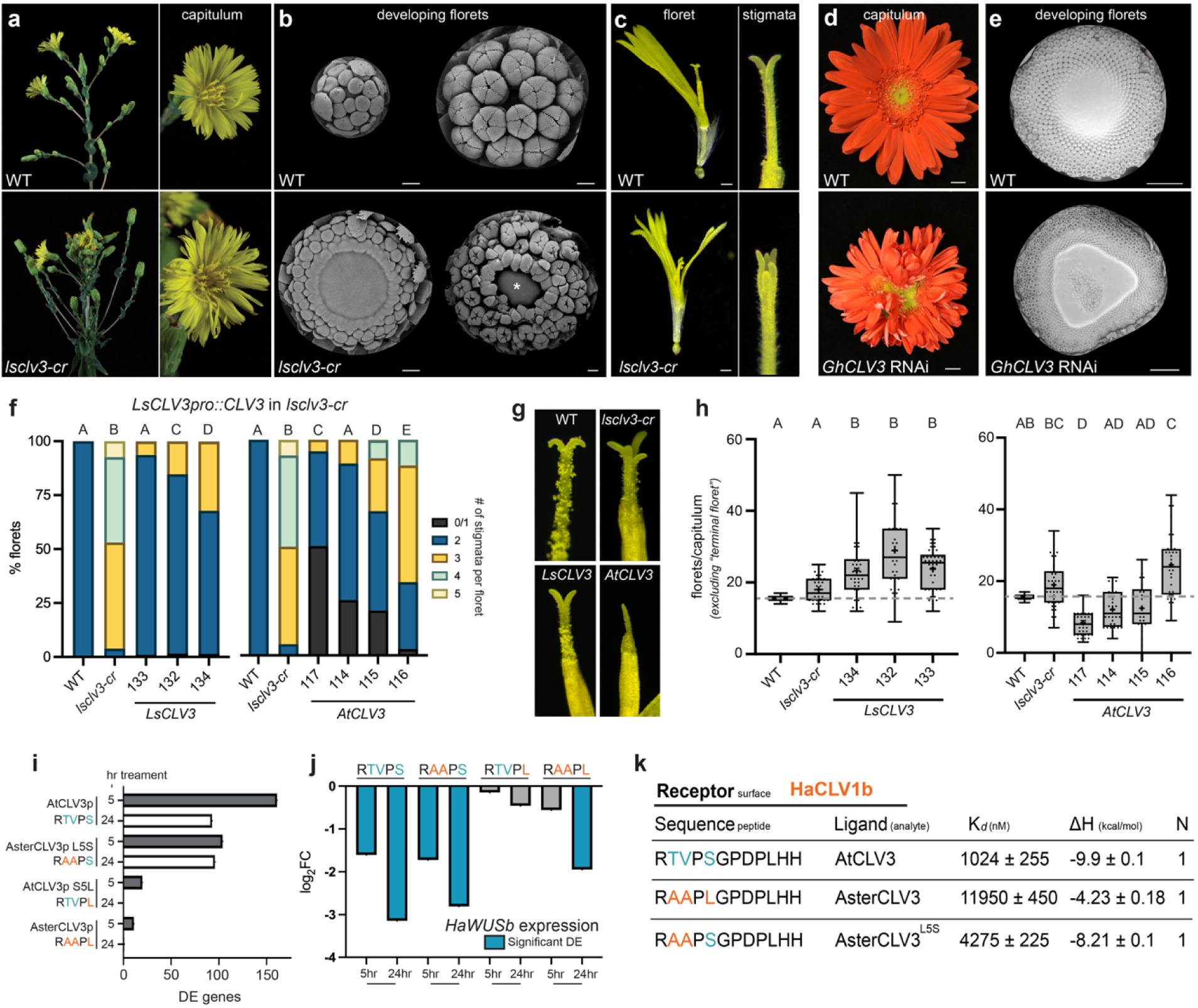
*AsterCLV3* has attenuated stem cell restrictive function. a-c, Capitula, scanning electron micrographs (SEM) of developing florets and mature florets of WT and *lsclv3-cr*. b, asterisk (*) indicates center of IM in *lsclv3-cr* with terminal floret developing. d-e, WT and *GhCLV3* RNAi knockdown capitula and SEM images of developing florets. f-g, Complementation of increased stigmata number phenotype comparing *LsCLV3pro::LsCLV3* (lines 133, 132 and 134) and *LsCLV3pro::AtCLV3* (lines 117, 114, 115 and 116) expressed in *lsclv3-cr*. Statistical summaries from Kruskal-Wallis and Dunn’s multiple comparison tests where A-E groups are significantly different (adj. p-value <0.0002). h, Floret counts per capitulum (excluding terminal florets) comparing *LsCLV3pro::LsCLV3* and *LsCLV3pro::AtCLV3* expressed in *lsclv3-cr*. Statistical summaries from Kruskal-Wallis and Dunn’s multiple comparison tests where A-D groups are significantly different (adj. p-value <0.05). (Related to Supplementary Fig. 17a-f). i-j, differential gene expression of sunflower inflorescence meristem cores 5 and 24 hours after treatment with CLV3p and CLV3p variants vs no peptide mock treatment. i, total differentially expressed genes (adj. p-value <0.05) (related to Supplementary Fig. 17). j, log_2_ fold change of *HaWUSb* expression after peptide treatment. k, ITC summary table of CLE/CLV peptides binding to the HaCLV1b ectodomain. K*_d_*(nM) represents binding affinity, ΔH (kcal/mol) indicates reaction enthalpy, and N denotes stoichiometry (n= 1 for a 1:1 interaction). Data are mean ± SD from at least two independent experiments, with undetected binding shown as n.d. (related to Supplementary Fig. 18). In f,h, (n=individual plants with 4 capitula counted each), in order left to right: n= 8 (WT), 10 (*lsclv3-cr*), 11 (*133*), 9 (*132*), 11 (*134*), 9 (WT), 10 (*lsclv3-cr*), 10 (*117*), 9 (*114*), 6 (*115*), 9 (*116*). Varying residues labeled in orange (if similar to AsterCLV3p) and blue (if similar to AtCLV3p). See complete statistical analyses and sample sizes in Supplementary File 8. Scale bar - 100μm (b), 1cm (d), and 1mm (c, e).

Next, we asked if the reduction in signaling capacity of AsterCLV3p relative to other species is important for shoot stem cell homeostasis in the Asteraceae. We synthesized a *LsCLV3* native promoter construct (∼3kb upstream and ∼1kb downstream of *LsCLV3*) and determined the promoter activity across lettuce meristems using a nuclear localized GFP (*LsCLV3pro::HistoneH2AX-GFP*) co-expressed with a plasma membrane marker (*AtUBIpro::PM-tdTomato*). We observed expression of *LsCLV3pro::H2AX-GFP* in the central zone of VMs, TMs, IMs, across independent lines; however, expression across developmental stages was inconsistent and often patchy, suggesting the existence of additional *LsCLV3* promoter regulatory elements (Supplementary Fig. 15a-b). In contrast, we observed consistent expression of the *LsCLV3pro* reporter in FMs throughout early floral initiation all the way to floral organ formation, including at carpel initiation (Supplementary Fig. 15c-d). Using the *LsCLV3pro,* we then attempted to complement *lsclv3-cr* with transgenes encoding different versions of the CLV3p sequence, *LsCLV3*pro::*LsCLV3* (RAAPL)*, LsCLV3*pro::*LsCLV3* containing the *AtCLV3* CLEp motif (RTVPS) or containing the *Scaevola aemula CLV3 (SaCLV3 –* Goodeniaceae) CLEp motif (RTAPL, differing from Asteraceae only at residue 2). As the *LsCLV3pro* was most consistently active during flower development, we quantified complementation using stigma number and number of florets produced per capitulum across three (*LsCLV3*) or four (*AtCLV3* and *SaCLV3*) independent stable transgenic lines. Both *LsCLV3* and *SaCLV3* complemented stigma and floret number at similar levels (Fig. 5f-h; Supplementary Fig. 16a-c). In contrast, lettuce expressing *AtCLV3* displayed gain-of-function effects, with reductions in the number of stigma per floret to 1 or even 0 in ∼25-50% of plants across lines (compared to the invariant 2 stigmata in wildtype plants), and a reduction in the number of florets per capitulum compared to wild-type lettuce in 3 out of 4 lines (Fig. 5f-h). Interestingly, we observed increases in the number of florets per capitula in *LsCLV3* complementation lines, possibly attributed to the partial suppression of the terminal floret phenotype (Supplementary Fig. 16d-f). Gain-of-function phenotypes in the *AtCLV3* complementation lines were not due to higher transgene expression levels and both *LsCLV3* and *AtCLV3* complementation lines had similar impacts on native *LsCLV3* and *LsWUSb* expression in developing florets of *lsclv3-cr* (Supplementary Fig. 16g-i). As such, normal stem cell homeostasis in Asteraceae requires attenuated CLV3p receptor affinity.

We then wanted to determine if CLV3p efficacy correlates with altered downstream *CLV*-signaling outputs in Asteraceae capitula. To determine this, we used RNAseq to assay transcriptional responses of sunflower capitula to treatment with either AsterCLV3p, AtCLV3p, and corresponding CLEp 5^th^ residue swaps (AsterCLV3p^Leu5Ser^ and AtCLV3p^Ser5Leu^), as our biological and genetic analysis demonstrates that the conserved AsterCLV3p Leu5 impairs signaling capacity and receptor binding. Sunflower was used because the larger inflorescence meristems of sunflowers are readily accessible (easily dissected) and both *HaCLV1a* and *HaCLV1b* paralogs retain the canonical LRR6 glycine that shapes the pocket required for CLEp binding, allowing the assessment of AsterCLV3p signaling without confounding impacts from *CLV1* alleles with potentially reduced CLEp binding. As the outer waxy cuticle of inflorescence meristems blocks CLEp diffusion (*33*), we devised a strategy in which we isolated tissue cores from the center of expansion phase sunflower capitula prior to floret differentiation (IM – 30d old sunflowers), incubated these IM tissue cores with CLV3p containing solutions and then compared transcriptional response to cores incubated with control solutions (Supplementary Fig. 17a-b). Tissues were collected for RNAseq 5 and 24 hrs post-treatment, a period that did not impact capitulum cell viability. Sunflower capitula responded most strongly to AtCLV3p (RTVPS), with 161 and 93 DEGs at 5 hrs and 24 hrs, respectively (Fig. 5i). Ser5Leu substitutions in AtCLV3p (RTVPL) reduced the number of DEGs, with 21 DEGs at 5 hrs and 0 at 24 hrs, consistent with a reduction in signaling capacity and the critical role for Ser5 in receptor binding (Fig. 5i). In contrast to AtCLV3p, sunflower capitula responses to their own AsterCLV3p (RAAPL) were strongly attenuated, with just 11 DEGs at 5 hrs, and only 1 DEG at 24 hrs (Fig. 5i). Capitulum transcriptional responses to AsterCLV3p were enhanced in AsterCLV3p^Leu5Ser^ (RAAPS) treatments, with 104 (5 hrs) and 96 (24 hrs) DEGs, again highlighting the importance of Ser5 for CLV3p signaling (Fig. 5i). We also observed overlap in transcriptional outputs coincident with Ser5 with AtCLV3p (RTVPS) and the AsterCLV3p var (RAAPS) at both 5 hrs (53 shared DEGs) and 24 hrs (54 shared DEGs) (Supplementary Fig. 17c-d, Supplementary Files 6-7). Remarkably, the 1 DEG identified in AsterCLV3p 24hr treatments was *HaWUSb*. In other Angiosperms studied, *WUS* is rapidly downregulated in response to CLV3p (*13*, *23*). *HaWUSb* was significantly downregulated at 5 hrs and 24 hrs by both AtCLV3p and AsterCLV3p^Leu5Ser^ (Fig. 5j). In contrast, AsterCLV3p treatments delayed *HaWUSb* downregulation as *HaWUSb* was only significantly repressed 24 hrs post-treatment. As such, the reduced capacity of AsterCLV3p to repress stem cell proliferation in Asteraceae capitula is associated with dampened downstream CLV transcriptional outputs, including the repression of *WUS*. To test the biochemical relevance of these results we successfully purified the ectodomain of sunflower CLV1b (HaCLV1b), and performed binding assays with AtCLV3p, AsterCLV3p, and AsterCLV3p^Leu5Ser^ (Supplementary Fig. 18). AtCLV3p bound to HaCLV1b with the highest affinity, with AsterCLV3p displaying ∼10 fold lower affinity, mirroring the enhanced potency of AtCLV3p to repress *WUSb* in sunflower IMs and stem cell proliferation in lettuce relative to AsterCLV3p (Fig. 5k). Again, Leu5Ser substitutions enhanced AsterCLV3p-HaCLV1b binding, mirroring the role for this residue on *WUSb* repression in sunflower IMs and in other assays (Fig. 5k). Proline hydroxyprolination did not improve the relative binding affinity of AsterCLV3p or abolish the affinity gap between the peptides, as AsterCLV3p consistently displayed lower affinity than AtCLV3p for both AtBAM1 and HaCLV1b, indicating that sequence differences, rather than hydroxylation state, are the primary determinants of their differential receptor-binding affinities (Supplementary Fig. 19). Additionally, we purified the LsCLV1a ectodomain, in which the LRR6 Gly that shapes the hydrophobic CLEp-binding pocket is replaced by Gln (Q196), a substitution expected to disrupt the geometry of this pocket. Consistent with this, LsCLV1a showed no detectable binding to either AsterCLV3p or AsterCLV3p^Leu5Ser^ (Supplementary Figs. 9a-d and 18b-c). Collectively, our data support a model wherein the downstream repression of *WUSb* and stem cell proliferation by CLV signaling in Asteraceae are conserved but dampened due to a quantitative reduction in CLV3p signaling capacity and receptor binding, thereby permitting the relaxed stem cell homeostasis critical for normal capitulum development.

We next traced the evolution of *CLV3* across the Asterales, specifically focusing on Asteraceae outgroups within MGCA clade, using newly generated chromosome-level assemblies and transcriptomic datasets for species in Campanulaceae (*P. grandiflorus*), Menyanthaceae (*Nymphoides indica*), Goodeniaceae (*S. aemula)*, and Calyceraceae (*Acicarpha procumbens*) (*26*). Strikingly, *A. procumbens* (Calyceraceae), which has a cephalioid inflorescence that also requires prolonged stem cell proliferation as in Asteraceae capitula (*17*), contains a single CLE1D1 *CLV3* homolog (*ApCLV3*) whose CLEp motif was identical to the Asteraceae CLV3p (RAAPL; Supplementary File 3). Thus, the evolution of AsterCLV3p RAAPL predates a Calyceraceae/Asteraceae split, suggesting that it evolved only once across the MGCA clade. In contrast Menyathaceae CLV3p (NiCLV3p, *N. indica*) contained an RRVPA N-terminal peptide sequence, while the Goodeniaceae (SaCLV3p, *S. aemula*) N-terminal CLV3p residues contained an RTAPL. Notably, in our Arabidopsis *clv3* complementation assays RTAPL variants of AtCLV3p display intermediate signaling capabilities compared to AtCLV3p and AsterCLV3p; while SaCLV3p and LsCLV3p both complement *lsclv3-cr* to similar levels (Figs. 3b-c, 5f-g and Supplementary Fig. 16a-c). Therefore, the evolution of head-like inflorescences in the MGCA clade appears to have been paralleled by a stepwise reduction in CLV3p activity and receptor affinity, from high activity in the Campanulaceae, to intermediate activity in the Goodeniaceae, to low affinity in the Calyceraceae and Asteraceae.

## Discussion

The re-wiring of conserved developmental networks is thought to underly evolutionary novelty in body plans across kingdoms, but few mechanistic examples outside of cis-regulatory re-wiring exist. Here we show that evolutionary innovation in Asteraceae inflorescence form was accompanied by quantitative reductions in CLV3p signaling due to a novel combination of substitutions in the AsterCLV3p CLE domain which lower signaling capacity and receptor affinity. At the same time, AsterCLV3p retains all essential receptor binding CLEp residues, thereby ensuring that affinity for CLV1/BAM receptors and signaling capacity is not eliminated entirely. This balanced signaling capacity correlates with reduced repression of *WUS*, a conserved target of CLV signaling in Angiosperms. The reduced signaling threshold promotes stem cell proliferation sufficient for normal capitulum development, while avoiding the adverse effects of unchecked proliferation. This is likely to explain the deep conservation of the peptide and the maintenance of at least one *CLV1* paralog with potential peptide-binding capability across the Asteraceae. Why some Asteraceae also evolved a putative dominant negative CLV1 paralog is not clear but generally mirrors the reduction of CLV3p signaling activity across the family. In Arabidopsis BAM and CLV1 receptors participate in CLV3-depedent shoot stem cell regulation, which is likely the case in other models as *clv1* mutants are generally phenotypically weaker than *clv3* mutants (*21*, *24*, *25*, *55*–*58*). While the relative contribution of BAM and CLV1 receptors to shoot stem cell regulation is not yet known in Asteraceae, or any other non-Arabidopsis model, our data suggests that AsterCLV3p substitutions impact binding and signaling through both BAM and CLV1 receptors classes. AsterCLV3p displays reduced affinity for HaCLV1b relative to AtCLV3p and undetectable binding to any BAM receptor we tested, consistent with an inability to promote *BAM1/2*-dependent ground tissue divisions in Arabidopsis, suggesting differences between BAM and CLV1 receptors in the perception of CLV3p. It is not yet clear if there is lower affinity for CLV1 from other species, but AsterCLV3p has intermediate signaling capability in Arabidopsis shoots suggesting this is likely the case. While our data support a model whereby AsterCLV3p substitutions impact signaling through reduced receptor affinity, we cannot fully rule out potential impacts on peptide diffusion or stability *in vivo*. Beyond shoot development, CLE peptides have diverse roles in plant growth. Our structural and biochemical data reveal distinct peptide-binding mechanisms that contribute to specificity in CLEp-receptor interactions. Notably, Ser5 plays an important role in modulating CLE peptide activity and receptor binding across different CLEp-receptor clades, which differs for the requirement of TDIF Ser5 in PXY interactions. Our findings provide a molecular mechanism for the meristem expansion central to both the *inflorescence-origin* and *floral-unit-meristem* models of capitulum evolution. The emergence of the RAAPL CLV3p predates Calyceraceae/Asteraceae divergence, with both families producing head-like inflorescences. Despite this, Calyceraceae are a small and geographically isolated family, while the Asteraceae spread globally from their ancestral origin to become the largest flowering plant family on earth (*26*). While our data suggest that reduced CLV3p signaling is important for normal capitulum development, lettuce *clv3* mutants and *AtCLV3* complemented lettuce *clv3* mutants still possess inflorescences patterned like heads. As such, other evolutionary innovations likely contributed to capitulum evolution and the success of Asteraceae, possibly through diversification of pollinator recruitment, seed dispersal traits, and the evolution of a rich arsenal of secondary metabolites (*1*, *10*, *59*). Moreover, recent findings suggest that ancient genome duplications, along with gene family expansions and subsequent selection, played a pivotal role in enabling Asteraceae to diversify and successfully disperse across the globe (*26*). Several data points suggest that the Goodeniaceae CLV3p ortholog is likely intermediate in signaling capacity between AsterCLV3p and outgroup CLV3p orthologs. Unlike Calyceraceae and Asteraceae, species in the Goodeniaceae exhibit variable inflorescence forms, with some possessing head-like inflorescences (*60*). This raises the possibility that variation in inflorescence architecture may be linked to intermediate CLV3p activity. Finally, our work suggests that engineering receptor-ligand affinity could provide a novel avenue for the quantitative re-shaping of crop stem cell regulation.

## Acknowledgments

We thank USDA GRIN, ABRC stock center, and Richard Michelmore for sharing seeds. We thank J. Mauricio Bonifacino for his plant collections and invaluable insights into capitulum morphology and evolution. We also thank Paige Ellestad for helpful discussions on gene family evolution. We thank Tony D. Perdue and Nathanaël Prunet, former and current directors of the UNC Biology Microscopy Core, as well as the Auburn University Research Instrumentation Facility and the Clemson Light Imaging Facility for assistance and access to imaging. We thank Jamie Winshell and James Garzoni for lab and plant growth facility support. We thank UNC’s High-Throughput Sequencing Facility for sequencing services.

## Funding

National Institutes of Health R35 MIRA grant R35GM119614 (Z.L.N).

National Science Foundation Plant Genome Research Program (PGRP) grant IOS-1546837 (Z.L.N.).

National Science Foundation Postdoctoral Research Fellowship in Biology grant PGRP IOS-1906389 (D.S.J.)

National Science Foundation Plant Genome Research Program (PGRP) grant IOS-2214474 (D.S.J.).

National Science Foundation Plant Genome Research Program (PGRP) grant IOS-2214473 (J.M.B.).

National Science Foundation Plant Genome Research Program (PGRP) grants IOS-2214472 and DBI-2319905 and DEB-1745197 (J.R.M).

Swiss National Science Foundation grant no. 310030_204526 (J.S.). University of Lausanne Fellowship (P.J-S).

Research Council of Finland project 341774 (P.E.).

Jane and Aatos Erkko Foundation (ASTROS: How meristems guide plant reproductive development? (P.E.).

## Author contributions

Conceptualization: ZLN, DSJ

Methodology: ZLN, DSJ, JS, JRB, JRM, PE, PJ-S

Investigation: ZLN, DSJ, RS, ADC, ACW, KRC, ATD, VG, RS, PJ-S, OR, FW, TZ, EMP, EY, JB.

Visualization: DSJ, ZLN, RS, ACW, PJ-S, KRC, JS, EY, JB, FW, TZ

Funding acquisition: ZLN, DSJ, JS, JRB, JRM, PE

Project administration: ZLN, DSJ

Supervision: ZLN, DSJ, JS, JRB, JRM, PE

Writing – original draft: ZLN, DSJ

Writing – review & editing: ZLN, DSJ, RS, JS, JRB, JRM, PE, PJ-S

## Data and materials availability

RNA-seq data from all Asteraceae species sequenced are available on the National Center for Biotechnology Information in BioProject PRJNA1261082. Sunflower data is in SAMN48412296, and all other species can be found in SAMN48412297-SAMN48412302. RNA-seq data from lettuce was obtained from Chen *et al*(*30*). All seed and vector materials are available upon request to the corresponding authors.

## Supplementary Materials

**Supplementary Table 1** – Oligos used for cloning, genotyping and *in situ* hybrization experiments.xlsx

**Supplementary File 1 –** Sunflower capitulum developmental transcriptomics.xlsx

**Supplementary File 2 –** genomes used for gene trees and ortholog identification.xlsx

**Supplementary File 3 –** CLE classification.xlsx

**Supplementary File 4 –** *CLV3* tblastn results – ortholog validation.xlsx

**Supplementary File 5 –** *CLV1* tblastn results – ortholog validation.xlsx

**Supplementary File 6 –** CLV3p treated sunflower inflorescence meristem RNAseq analysis – 5hr.xlsx

**Supplementary File 7 –** CLV3p treated sunflower inflorescence meristem RNAseq analysis – 24hr.xlsx

**Supplementary File 8 –** Statistical analyses used throughout study.xlsx

**Supplementary File 9 -** CLV1/BAM receptors and WUS gene trees (higher resolution)

**Supplementary File 10 -** CLV1 conserved binding pocket alignment (higher resolution)

**Supplementary File 11 -** BAM1 and CLV1 share a conserved peptide-binding pocket (higher resolution)

## Materials and Methods

### Plant material and growth conditions

#### Sunflower, lettuce, Gerbera and additional Asteraceae species accessions

*Helianthus annuus* cv Ha412HO (sunflower; PI 642777), *Lactuca sativa* (lettuce; PI 251246), *Calendula suffruticosa* (PI 607419), *Gaillardia pinnatifida* (W6 55897), *Packera multilobata* (W6 30293), *Stokesia laevis* (PI 346981), and *Symphyotrichum novae-ang*liae (PI 667514) were all sourced from the USDA National Plant Germplasm System. *Bidens ferulifolia* cv. Compact Yellow was acquired as cuttings from North Carolina Farms Inc. Micropropagated *Gerbera hybrida* cv. Terra Regina plantlets were provided by Dümmen Orange, The Netherlands. *H. annuus* cv. Pacino cola seeds were purchased from Siemenliike Siren Oy, Finland.

#### Asteraceae growth conditions

Sunflower achenes were sown on pre-wet vermiculite (A-3 coarse) and germinated under a dome at the UNC greenhouse in long-day conditions; LD, 16hr light/8hr dark (light supplemented to 400W/m^2^) with temperature kept between 20-25°C. At 7 days (7d) post germination, seedlings were transplanted to 4.5-inch geranium pots (Greenhouse Megastore CN-STD-0450) in soil (Metro-Mix 360/sand/perlite supplemented with Marathon pesticide) and continuously watered with Peter’s 20:20:20 [N:P:K] at 150ppm (parts per million). At 20d post germination, plants were then up-potted to 1-gallon pots in fresh soil and Osmocote [14:14:14] was added for top watering. Lettuce achenes were sown on 0.8% phytoagar with half-strength Murashige-Skoog (½ MS) media buffered with MES at pH 5.7 and germinated in a growth chamber (Percival CU-36L4) in LD conditions at 22°C for 5 d. Seedlings were transplanted to soil (2.8 cf PRO-MIX BX, 40g Marathon 1% G Insecticide, and 1 gal 200ppm Peter’s 20-10-20 fertilizer) in 6 cell seedling inserts (BWI FG12067) and grown in growth chambers (either Percival AR-66L2_or AR-75L2) in LD conditions at 22-25°C. 2-week-old seedlings were transplanted to 4.5 inch geranium pots (Greenhouse Megastore CN-STD-0450). 4-week-old plants were supported with bamboo stakes and thereafter fertilized weekly with 200 ppm Peter’s 20-10-20 fertilizer. Gerbera plants were all grown in a greenhouse as described previously (*61*). All other Asteraceae species were grown similar to sunflower methods above but in a custom-built grow room at UNC with environmental control (temperature maintained at 21-25°C) under red and blue LED lights in LD conditions.

#### Arabidopsis lines and growth conditions

Mutant alleles used in this study were all in the Col-0 (Columbia-0) background with specific allele information as follows: *clv3* (*clv3-9*) (*24*), *clv1* (*clv1-101*) (*14*), *bam1* (*bam1-4*) (*24*), and *bam2* (*bam2-4*) (*24*). The published endodermal reporter *EN7pro::YFP-H2B* (*50*) was used in root meristem analyses. All Arabidopsis were grown as follows: seeds were sterilized and plated on ½ MS media (with MES buffer at pH 5.7) and stratified in the dark at 4°C for 2 days. Plates were then either moved to a custom-built grow room at UNC with environmental control (temperature maintained at 21-25°C) or two Percival growth chambers (AR-75L3 at UNC or Auburn). By 10d, seedlings were transplanted to soil (Metro-Mix 360/sand/perlite supplemented with Marathon pesticide and Peter’s 20:20:20 at recommended levels) then grown in same conditions they were germinated in until mature and flowering for phenotypic analyses.

### Confocal microscopy

For sunflower, shoot apices from 10d, 20d, 30d, 35d, and 37d (days post germination) plants were removed, cut longitudinally and fixed overnight in a 1:20 (tissue:fixative) volume of FAA (2% formaldehyde, 5% acetic acid, 60% ethanol v/v) at 4°C with constant agitation on a rocker. Meristems were cleared and autofluorescence was imaged on a confocal laser scanning microscope as described previously for the 10d, 20d and 30d stages on a Zeiss 710 using a Plan-APOCHROMAT 10x objective (NA=0.45) (*21*, *62*). For 35d and 37d stages, meristems were imaged on a Zeiss 880 using an EC Plan-Neofluar 10x objective (NA=0.30) in tiling mode. For both stages, 24 images were stitched together using Nikon NIS-Elements to produce larger spatially resolved micrographs of entire developing capitula. For *Gerbera,* images of *proGhCLV3::NLSeGFP::GhCLV3ter* were taken with a Leica SP8 upright laser scanning confocal microscope equipped with a HC Fluotar L 25x/0.95 VisIR water dipping objective. The GFP was excited with a 488nm laser, and the emission was collected from 495nm to 545nm. Given the large size of the IM/FM stage, 9 tile scans were taken with a single sample and merged by the LAS X software. The images were then exported into tiff image stacks and further visualized with the MorphoGraphX software (*63*). For *Arabidopsis,* dissected root meristems of *EN7pro::YFP-H2B* were imaged on a Zeiss 710 using a C-Apochromat 40x water immersion objective (NA=1.20). Seedlings were fixed in room temperature 4% PFA for 4 minutes to preserve fluorophores. Fixed seedlings were stained with 0.1% Calcofluor white (Fluorescent Brightener 28; Sigma-Aldritch) in Clearsee (10% xylitol, 15% sodium deoxycholate, 25% urea) (*64*) and following dissection of the root tips mounted in Clearsee. Root meristems were imaged with following excitation/emission settings: YFP (514nm; 519 to 564nm) and Calcofluor white (405nm; 410 to 551nm). Unless specified, all images were processed as separate channels and then merged when necessary in Fiji (v.2.14.0/1.54f) (*65*). For lettuce, the *LsCLV3* transcriptional reporter was imaged on an inverted Leica Stellaris 5 using a Leica HC PL APO 20x/0.75 IMM CORR CS2 M25. Developing bracts and leaves were removed before shoot apices were cut longitudinally and placed in 50mM PIPES buffer in glass-bottom petri dishes (catalog no. P35G-1.5-10-C; MatTek Corporation). Coverslips were used to keep sample cut faces flat against the glass bottom. Nuclei were labeled using *LsCLV3pro::H2AXa-GFP* (488nm; 495-545nm). The first 8aa of SlCPK1 was translationally fused to the N-terminus of tdTomato and expressed under *AtUBIpro* to constitutively label plasma membranes (561nm; 575-600nm) (*66*).

### Scanning electron microscopy

For Arabidopsis, soil-grown plants with young inflorescences (4-5 cm in height) were selected for SEM imaging and IM measurements. Floral buds were carefully removed, and IMs were collected in ice-cold 100% methanol and gradually replaced with 100% ethanol before critical point drying (CPD). CPD was performed according to the manufacturer’s protocol (Leica EM CPD300). Scanning electron microscopy images were taken using a Hitachi TM4000Plus II SEM system, as described earlier in our lab (*35*). After taking the SEM images, IM size was measured using Fiji (https://imagej.net/software/fiji/downloads) software.

Lettuce inflorescences (WT and *lsclv3-cr*) were dissected and fixed in Formaldehyde Acetic Acid overnight, before serial dehydration in increasing concentrations of ethanol (70%, 80%, 90%,100%; 2hrs each step). The dehydrated samples were then critical point dried in Tousimis Samdri-PVT-3D and gold sputter coated using a Quorum EMS 150R ES. The samples were then imaged using a Zeiss EVO50. *Gerbera* inflorescences (WT and *GhCLV3* RNAi lines) were dissected, fixed and imaged as described previously (*67*) using a Quanta 250 SEM (FEI Corporation).

### Photography and stereoscope imaging

Images of lettuce stigmata, florets, and capitulescence meristems were captured on a stereo microscope (Leica M60, Flexacam C3). For *lsclv3-cr* IM and fasciation, the leaves of 38 day-old plants were removed to allow for a clear view of the primary stem while all branches were left intact. Images of shoot apices and meristems were taken on a Leica M165 FC stereo microscope with a plan apo 1.0x objective and a K3C camera on a 0.5x C-Mount. Photographs of inflorescences from *Arabidopsis*, lettuce and *Gerbera* were taken using a DSLR Camera (Cannon EOS Rebel T5 with a Tokina 100mm f/2.8 AT-X M100 AF Pro D macro lens or similar). Images were edited in Gimp (v2.10) or Adobe Photoshop. *Arabidopsis bam1/2* rosette phenotypes and carpel pictures were made using a Samsung Galaxy S23 phone. For *bam1/2* inflorescence pictures (sterility phenotype), source images were photographed using a Canon Eos Rebel T6i DSLR camera on a tripod equipped with a Canon EF-S 35mm f/2.8 Macro IS STM lens. A focus stack of varying focal planes through the sample were acquired, and composite inflorescence images were made using Affinity Photo (version 1.10.6.1665) Focus Merge tool.

### RNA sequencing and analysis

#### Staging, CLEp treatments and RNA extraction

For sunflower developmental RNAseq, shoot meristems from 10d (vegetative meristems), 20d (floral transition meristems), 30d (inflorescence meristems; IMs) and 35d (inflorescence and floral meristems) post-germination sunflower plants were dissected and flash frozen in 1.5mL microcentrifuge tubes in liquid nitrogen. Meristems were collected in three biological replicates with 10-20 meristems pooled per sample and stored at -80°C until further processing. For *de novo* transcriptome assemblies of Asteraceae species, shoot meristems were dissected spanning different stages of development including vegetative and floral (when available). Approximately 8-10 meristems were pooled into one sample per species, flash frozen in liquid nitrogen and stored at -80°C until processing. For both the sunflower developmental series and Aster shoot meristem transcriptomes, samples were hand ground into fine powder using small pestles fitted to 1.5mL microcentrifuge tubes in liquid nitrogen. For CLV3p variant IM response experiment, 30d sunflower shoot meristems were dissected and stored on inflorescence meristem media (1/2 MS minimal, 4% sucrose (w/v), 0.05% casein hydrolysate (w/v), pH 5.5, 100mg/L myo-inositol, 1.2mg/L kinetin, 0.1% plant preservation mixture (PPM), 1% agarose) while all ∼100 meristems for a single rep were dissected (∼1-2hrs). A single tissue core was taken from the center of each 30d IM using a 1.5mm Biopsy Punch (Miltex) to facilitate CLEp delivery to meristematic cells upon treatment. Approximately 8-10 30d IM cores were used for each treatment and replicate. Cores were incubated in inflorescence meristem media (without agarose) containing 0.1uM CLEp vars (no peptide control, AtCLV3p, AsterCLV3p, AtCLV3p S5L, and AsterCLV3p L5S) for 5hrs or 24hrs at room temp with constant agitation on a table-top plate rotator. Samples were then collected in 2mL microcentrifuge tubes and flash frozen in liquid nitrogen. Three biological replicates at 5hrs and 24hrs (30 total samples) were then ground using a liquid nitrogen cooled genogrinder (Spex). RNA was extracted from each sample using the EZNA Plant RNA kit (Omega Bio-tek) and treated with RNase-free DNase (Omega Bio-tek). Approximately 1.5μg of RNA was used in library preparation, using the stranded mRNA-seq kit (Kapa Biosystems) at the High-Throughput Sequencing facility at UNC. 50bp paired-end reads were generated on either an Illumina HiSeq 4000 or 6000 with read depths that varied across experiments. Samples for *de novo* transcriptomes were sequenced at ∼80-100 million (M) reads per sample, while samples for differential expression analyses were sequenced at ∼50-100M reads (developmental series) and ∼30-40M reads (CLEp treated IMs) per replicate.

#### Gene expression analyses

For sunflower, Illumina paired-end RNAseq reads were quality assessed (FastQC v0.11.9 (*68*), MultiQC v1.14 (*69*)) and genome indexed (STAR v2.7.10b (*70*)). Reads were mapped to the HA412-HOv2 sunflower reference genome (*19*) and raw read counts per gene were quantified (STAR v2.7.10b). Combat-Seq was used to reduce unwanted variation between biological replicates (sva v3.42.0 (*71*)). Differential gene expression analysis was done with DESeq2 (*72*) (v1.34.0) with the model ∼0 + development stage using a Wald test for significance. P-values were then corrected for multiple testing using the Benjamini-Hochberg procedure and only genes with an adjusted p-value (FDR) < 0.05 were considered significant. For clustering, per the DESeq Workflow, the count data was transformed using variance stabilizing transformation without blind dispersion estimation. Hierarchical clustering was done on the VST-transformed data in R (*73*) (v4.1.2) using hclust. All genes that were differentially expressed between the pairwise comparisons are included (12,657 genes total). The distance matrix was calculated with Euclidean distance and the complete linkage method for clustering was used. The tree was cut at a height of 2.7 (Supplementary Fig. 1b). For lettuce, publicly available datasets were downloaded using sratoolkit (data: SRR6388382-SRR6388389 (*30*)) and paired-end reads were mapped to the lettuce reference genome (v8 (*74*)) using hisat2 (*75*) (v2.2.1). Stringtie (*76*) (v2.1.1) was used to produce a shoot meristem transcriptome-based annotation (*LsCLV3* is unannotated in the available reference genome annotation) which was then leveraged to quantify gene-level expression of the stem cell regulators *LsCLV3* and *LsWUSb* using featureCounts in subread (*77*).

#### *De novo* transcriptome assembly

Illumina paired-end RNAseq reads from shoot meristems (SAMs) of six Asteraceae species (Supplementary Fig. 2c) were quality assessed (FastQC v0.11.9 (*68*)) then assembled into *de novo* transcriptome assemblies using the Trinity pipeline (*78*, *79*) (v.2.11.0) containing SAM-expressed genes. Coding regions of transcripts were identified and protein sequences extracted using the TransDecoder (*80*) (v5.5.0) LongOrf and Predict functions with -m flag set at 60 to capture smaller proteins of potential interest (*i.e* CLE peptides). The protein fasta file output from each Asteraceae SAM transcriptome were used for downstream identification of stem cell signaling orthologs.

### Ortholog Identification

#### Proteome analysis for CLV3 and CLV1 across Angiosperms

To identify CLV1 and CLV3 sequences in Asteraceae, we analyzed a comprehensive set of complete proteomes from 13 Asteraceae species and 37 outgroup species, representing major orders from Monocots, Magnoliids, and the ANA grade (taxonomic information and accession numbers are available in Supplementary Fig. 2 and Supplementary File 2). Among these, eight represent newly assembled genomes from the Asterales (Moore-Pollard *et al.* 2025), and our analysis of *CLV1*-related genes was extended to include the six unpublished *de novo* transcriptomes (derived from shoot meristems; see above) from diverse Asteraceae species.

#### CLE-clustering and identification of AsterCLV3

For the *CLV3* analysis, we conducted two independent HMMER3 (*81*) screenings using profiles of CLE motifs derived from Oelkers et al. (2008) (*82*) and Zhang et al. (2020) (*28*), applying a domain e-value threshold of 40 to reduce the chance of false positives. Since CLE peptides are unsuitable for standard phylogenetic methods due to their small size and high sequence variability outside the conserved domain, we employed a phenetic clustering approach using CLANS (*83*) to identify putative Asteraceae orthologs of *CLV3*. During the initial run, we tested seven e-value thresholds for high-scoring segment BLASTP (*84*) pairs, ranging from 1e-04 (default) to 1e-10. The screening based on the Zhang et al. profile was slightly more conservative (retrieving 1052 sequences vs. 1200), resulting in fewer singletons; thus, we used this dataset for the final analyses. An e-value threshold of 1e-7 produced clustering patterns consistent with previous studies (*27*, *82*), including a clearly defined 1D1 cluster with canonical *CLV3* from *A. thaliana*.

#### CLV1 Ortholog Identification

The identification of CLV1 genes involved BLASTP screening, followed by InterProScan (*85*) scanning to confirm the presence of the signature protein-kinase domain (PF00069) and leucine-rich repeat (LRR) regions (PF00560) at an e-value of 1e-10. In transcriptomes where multiple splicing variants were represented, we retained the longest variant for subsequent analyses. The initial search retrieved 1065 genes, of which 302 were confirmed as belonging to the CLV1/BAM family based on preliminary phylogenetic inference. For the final reconstruction, sequences were re-aligned using the G-INS-i algorithm implemented in MAFFT (*86*) (v7.271). After manually correcting misaligned positions, we collapsed duplicated sequences using ElimDupes and performed quality trimming with trimAl (*87*) (v1.2rev59). Lastly, phylogeny was reconstructed using RaxML (*88*) (v8.2.4), with automatic protein model selection (-m PROTGAMMAAUTO flag), and branch support estimated with 200 rapid bootstrap replicates.

CLV1a, CLV1b, and CLV3 from *Helianthus annuus* L. and *Lactuca sativa* L. were queried against coding sequences of eight Asterales species (with reference-level assemblies and new gene annotations) (*26*) using TBLASTN (*89*) (v2.13.0). CLV genes of each taxon were confirmed by having the lowest e-value and highest score from BLAST (Supplementary Files 4-5). *GhCLV1a* and *GhCLV1b* were identified from Gerbera transcriptome data sets and the sequences were amplified and validated by sequencing.

### *In situ* hybridization

The cultivars used for *in situ* hybridization are Gerbera cv. Terra Regina, sunflower cv. Pacino cola, and lettuce (PI 251246). *GhWUSb*, *HaCLV1a* and *LsCLV1a in situ* hybridization was done with gene-specific probes, and *GhCLV1a*, *GhCLV1b*, *HaCLV1b*, *HaWUSb*, *LsCLV1b*, and *LsWUSb* with full length hydrolyzed probes. *CLV3* genes were done with full length probes. The probes were amplified with PCR. After purification, the probes were synthesized as previously described (*90*) and labeled with DIG RNA Labeling Kit (Roche) according to the instructions. When appropriate, the labeled probes were further hydrolyzed to approximately 150bp (*91*). The preparation of plant samples, sectioning, and hybridization were done as described by Elomaa et al. 2003 (*92*). Oligo sequences used for *in situ* hybridization are found in Supplementary Table 1.

### Root CLE peptide treatments

All CLE peptides and variants used in this study were synthesized by Biomatik (>90% purity) as in previous studies (*49*).

#### CLE peptide variant root length assay

Arabidopsis (Col-0) seeds were sterilized then plated on a fine mesh placed on ½ strength Murashige and Skoog medium supplemented with 0.8% Phytoagar. The seeds were stratified for 48 hours at 4°C, then germinated vertically under 24-hour light at 22°C for four days. The fine mesh that the seedlings were germinated on was then transferred to ½ strength Murashige and Skoog medium supplemented with 0.8% Phytoagar and 0.1μM mock treatment, CLEp or CLEp variants (with S5L substituions) under sterile conditions. Seedlings grew for 4 days vertically following the transfer, then plates were scanned and root lengths were measured using Fiji (*65*).

#### CLV3p root length assay

For Arabidopsis, seeds were sterilized then plated on ½ strength Murashige and Skoog medium supplemented with 0.8% Phytoagar, and either 0.1μM of CLV3p (from Arabidopsis, Asteraceae, outgroup taxa or amino acid substitutions as indicated) or varying concentrations of Arabidopsis and Asteraceae CLV3p (0μM, 0.01μM, 0.1μM, 1μM, and 10μM). The seeds were stratified for 48 hours at 4°C, then germinated vertically under 24-hour light at 22°C for ten days. Plates were then scanned, and root lengths were measured using Fiji (*65*). For lettuce and sunflower, achenes were sterilized and plated on ½ MS buffered with MES at pH 5.7 containing 0.1μM of CLV3p (species and amino acid substitutions as indicated). For sunflower, achenes were grown in sterile cups containing ∼200mL of media for ten days in long-day conditions. Only seedlings with fully expanded cotyledons and true leaves initiated were used for root length comparisons. For lettuce, achenes were grown vertically on plates in 24-hour light at 22°C for seven days. Plates were scanned and primary root length measured using Fiji (*65*).

### Transgenic lines

For *Arabidopsis,* new transgenic lines were generated through floral-dip transformation (*93*) of binary vectors into indicated backgrounds using *Agrobacterium tumefaciens* (GV3101). Transgenic plants were selected for on ½ MS plates using appropriate antibiotic selection (Kanamycin or Hygromycin). Unless specified differently, T1 plant lines were used to assess transgene function across many (typically >10) independent transgene insertion events to reduce selection bias.

For lettuce, achenes were germinated as described above. 5-day-old cotyledons were removed, and their apical tips were dissected off with a razor blade. Cotyledon explants were added to a suspension of *A. tumefaciens* (GV3101) and nutated for 5 min. *A. tumefaciens* inoculation suspension was created by pelleting 2 mL of overnight culture and resuspending in 20 mL inoculation solution (½ MS salts and vitamins, 1% sucrose, 0.5 g/L casamino acids, 200 mM acetosyringone, pH 5.33). Explants were lightly patted dry to remove excess *A. tumefaciens* and placed abaxial side down on co-cultivation media (½ MS salts, full strength MS vitamins, 3% sucrose, pH 5.7, 0.8% phytoagar, 0.1 mg/L NAA, 0.5 mg/L BAP, and 200 mM acetosyringone). A piece of sterile filter paper was placed between explants and co-cultivation media to prevent *A. tumefaciens* overgrowth. Co-cultivation plates were then kept in the dark for 3 days at 22°C. Explants were then transferred to callus induction media containing Timentin (RPI T36000-5.0) to kill *A. tumefaciens* and Kanamycin for antibiotic selection (½ MS salts, full strength MS vitamins, 3% sucrose, pH 5.7, 0.8% phytoagar, 150 mg/L Timentin, 100 mg/L kanamycin, 0.1 mg/L NAA, and 0.1 mg/L BAP) and kept at 22°C for 2 weeks in a tissue culture chamber (Percival CU-36L4). The resulting callus and nascent shoots were then transferred to shoot elongation media (½ MS salts, full strength MS vitamins, 3% sucrose, pH 5.7, 0.8% phytoagar, 150 mg/L Timentin, 100 mg/L kanamycin, and 0.01 mg/L BAP) for 1-3 weeks. Once shoots elongated, they were excised from surrounding callus and placed in rooting media (½ MS salts, full strength MS vitamins, pH 5.7, 0.8% phytoagar, 150 mg/L Timentin, 100 mg/L kanamycin, and 0.1 mg/L NAA). Once roots developed, plants were transferred to soil, covered with humidity domes for 2 d before acclimated to chamber conditions fully. For Gerbera, wild-type *Gerbera hybrida* cultivar ’Terra Regina’ transformation was performed as previously described (*94*).

### Molecular cloning

Primer sequences used for cloning can be found in Supplementary Table 1.

For the generation of *CLV1* and *BAM1* mutant vectors, recombinant PCR was used to create LRR DNA fragments with point mutations in *BAM1* or *CLV1* as described before (*24*). Fragments were cloned into the pBLUNTII Topo vector (Invitrogen) and sequenced. Fragments were then subcloned into *pENTR* vectors containing the corresponding wild-type full length genomic-2XGFP receptor clones, sequenced, and then recombined into previously created gateway compatible native promoter binary vectors (*24*). For *CLV3* variants, *CLV3* variants were synthesized by Twist Bioscience into manufacturer provided entry plasmids and then recombined into a previously created gateway compatible native promoter *CLV3* binary vector.

Entry plasmids (*pENTR*) containing the *LsCLV3* (lettuce) genomic sequence with CLE domain encoding different variants (*AtCLV3, AsterCLV3, and SaCLV3*) and an empty vector containing both 5’ upstream and 3’ downstream *LsCLV3* cis-promoter regions were synthesized by Twist Bioscience. *LsCLV3* variants, as well as *H2AX-GFP*, were amplified by PCR using cloning primers with PacI sites, then digested and ligated into the *LsCLV3pro* binary to form expression vectors. All binary vectors were introduced into chemically competent *Agrobacterium tumefaciens* (GV3101).

*GhCLV3* full length coding sequence was cloned into the Gateway compatible RNAi vector *pK7GWIWG2*(II) (*95*). For *GhCLV3* transcriptional reporter construct, 1652bp upstream and 236bp downstream of *GhCLV3* were cloned into entry vectors *pDONR*_*Amp*_*P4P1R* and *pGEMtP2rP3* as promoter and 3’ sequences (both entry vectors were kindly provided by Prof. Ari Pekka Mähönen, University of Helsinki, Finland). The sequences were verified by plasmid sequencing. An LR reaction was performed to clone the promoter, *NLSeGFP* and 3’ sequences into destination vector *pKm43GW*(*95*). The constructs were transferred into the *A. tumefaciens* strain C58C1.

### Lettuce mutant analyses and Gerbera RNAi

For lettuce, two *lsclv3* mutant alleles were generated, a +1 insertion (*lsclv3-cr*) and a -6 deletion (*lsclv3-cr2*) (Supplementary Fig. 14a). *lsclv3-cr* was detected by dCAPS PCR and BslI digestion, and *lsclv3-cr2* was detected by PCR amplification and MwoI digestion with listed primers (Supplementary Table 1). As *lsclv3-cr* plants rarely produced viable seed, and transgenic *LsCLV3* variants weren’t easily distinguished from WT *LsCLV3* by PCR, transgenes were introduced into an *lsclv3-cr/lsclv3-cr2* biallelic mutant background (as the *lsclv3-cr2* -6 deletion is in frame and there was no detectable phenotype). Transgenic achenes were sown on ½ MS media with kanamycin for antibiotic selection and confirmed using a *AtUBI10::tdTomato* selectable marker. All complementation lines were confirmed as both transgene positive and *lsclv3-cr* -/- mutants prior to phenotypic analysis.

For *Gerbera,* six independent transgenic GhCLV3 RNAi lines showed fasciated inflorescence phenotypes. The transgenic line TR5 was selected for gene expression analysis. For wild-type plants the undifferentiated meristem was dissected from inflorescences of 2 to 3 mm in diameter. For *GhCLV3* RNAi plants only fasciated undifferentiated meristem was dissected. Three independent biological replicates, each with at least three meristem samples, were collected for transgenic and wild-type plants. Total RNA preparation, cDNA synthesis (500 ng total RNA was used), quantitative RT-PCR, and data analysis were done as previously described (*96*). *GhCLV3* and *GhWUSb* results were compared with a Welch’s t-test.

### RT-qPCR analyses

For Arabidopsis, total RNA was isolated from 100 mg of seedling tissue from 9-day-old plants grown on ½ MS plates. 3-4 biological replicates were used for RNA isolation. RNA was isolated using RNAzolRT (Sigma, R4533-50ML), 4-bromoanisole (MRC, BN 191), and Glycoblue (Thermo, AM9515) as per the manufacturer’s (Sigma) protocol. First-strand cDNA synthesis was performed using the Protoscript II First Strand cDNA Synthesis Kit (NEB, E6560L) as earlier described (*97*). RT-qPCR was performed using PowerUp SYBR Green Master Mix (qPCRBIO SyGreen Blue Mix Lo-ROX, Genesee 17-505B) and the Applied Biosystems QuantStudio 6 Flex System. *PP2A* transcripts as the internal control (*97*). Due to the high degree of sequence similarity, we designed and used the *CLV3* RT-qPCR primers, which can detect and measure the transcripts from both the endogenous and the transgene *CLV3*. RT-PCR was done using cDNA as a template with Red Taq 2X Master Mix (VWR, 76620-472). Primer information is included in Supp Table 1*. AtCLV3* relative expression was analyzed using a mixed model with genotype as a fixed effect and independent transgenic lines as a random effect, fit by REML (lme4 2.0.1) (*98*), with Satterwaite degrees of freedom (lmerTest 3.2.1) (*99*). All pairwise comparisons were tested jointly using a single-step multivariate-t adjustment (emmeans 2.0.3) (*100*). To assess sensitivity to deviations from normality, the model was refit using a robust model (robustlmm 3.5.0.2) (*101*); Results were consistent between models.

For lettuce, developing lettuce florets were collected from capitula terminating secondary branches, similar to those used for stigma counts. Each sample comprised florets from 8-12 capitula from 2-3 individuals. RNA was isolated using E.Z.N.A.® Plant RNA Kit (Omega Bio-tek #R6827) with an on column DNase digestion (Omega Bio-tek #E1091). Samples were run on a gel to check for degradation and DNA contamination before cDNA synthesis using a ProtoScript® II cDNA synthesis kit (NEB #E6560) with oligo-dT primers. Samples with the highest RNA concentrations were used for NRT controls and yielded no qPCR product. The cDNA equivalent of 2ng of RNA was used in qPCR reactions with Luna® Universal qPCR Master Mix (NEB #M3003). *LsUBI10* was used as a reference gene with primers spanning the intron (285bp amplicon). All *LsCLV3* primers spanned both introns. The primers used to amplify endogenous *LsCLV3* also amplified transgenic *LsCLV3* due to high sequence similarity (293bp amplicon). The 3’ end of the forward primer used to detect only transgenic *LsCLV3* annealed to a PacI cloning scar immediately upstream of the start codon absent from endogenous transcripts (327bp amplicon)—control reactions using WT and *lsclv3-cr* samples yield no product. *LsWUSb* primers spanned 2 introns (261bp amplicon). Lettuce RT-qPCR reactions were run on a BioRad CFX Connect. NTC reactions yielded no product; samples with the highest RNA concentrations were used for NRT controls and yielded no qPCR product. Because 3-4 independent lines do not justify treating transgene insertion as a random effect, relative expression of *LsWUSb* and total *LsCLV3* was instead modeled using generalized least squares (nlme 3.1.169) (*102*) allowing group-specific residual variance (varIdent) to account for heteroscedasticity. To avoid undue multiplicity burden in treating identical transgenes as independent, nine pairwise contrasts—each sample compared to WT and *lsclv3-cr*—were made using the same single-step multivariate-t adjustment (glht; multcomp 1.4.30) (*103*). *LsCLV3* transgene levels were compared with a Welch’s ANOVA and a Games-Howell post-hoc test.

For *Gerbera,* six independent transgenic *GhCLV3* RNAi lines showed fasciated inflorescence phenotypes. The transgenic line TR5 was selected for gene expression analysis. The undifferentiated meristem was dissected from inflorescences of 2 to 3 mm in diameter from both *GhCLV3* RNAi and wild-type plants. Three independent biological replicates, each with at least three meristem samples, were collected for transgenic and wild-type plants. Total RNA preparation, cDNA synthesis (500 ng total RNA was used), quantitative RT-PCR, and data analysis were done as previously described (*96*). *GhCLV3* and *GhWUSb* results were compared with a Welch’s t-test.

All RT-qPCR analyses performed on ΔCq values and plotted as 2^-(ΔCq)^. Mean values and SEM limits were calculated using ΔCq values then plotted as 2^-(ΔCq)^. All analysis scripts and processed data are available on GitHub (https://github.com/thecapitulab/AsterCLV3-qPCR-and-carpel-models) and archived at Zenodo (https://doi.org/10.5281/zenodo.22184594). Analyses were performed in R 4.6.0.

### Phenotypic analyses

All phenotypic data was collected and analyzed using Prism 10 (GraphPad), unless stated otherwise below. Details of statistical analyses can be found in Supplementary File 8.

Leaf numbers for the Arabidopsis *clv3* complementation experiment were made on 3-week-old soil grown plants using a tabletop dissecting scope. Stem width measurements were performed in the same experiment as described before (*35*). *bam1 bam2* rosette phenotypes were scored in 3-week-old soil grown plants, and fertility was confirmed by the production of siliques containing seeds. All Arabidopsis carpel number quantifications were made as described previously (*35*), with 15 flowers counted per T1 plant (*clv3* experiments), and 20 flowers counted per T1 (*clv1* experiments). *bam1 bam2* inflorescences were photographed using a Canon Eos Rebel T6i DSLR camera on a tripod equipped with a Canon EF-S 35mm f/2.8 Macro IS STM lens. A focus stack of varying focal planes through the sample were acquired, and composite inflorescence images were made using Affinity Photo (version 1.10.6.1665) Focus Merge tool.

Arabidopsis carpel counts were modeled using a COM-Poisson generalized linear mixed model via compois (glmmTMB 1.1.14) (*104*) with genotype as a fixed effect and independent transgenic line as a random effect. This model was chosen over a standard Poisson GLMM due to significant underdispersion, exemplified well by the highly canalized development of 2 carpels in Col-0. COM-Poisson GLMMs introduce φ as a parameter for dispersion, while φ is functionally fixed at 1 in standard Poisson GLMMs. φ was allowed to vary by genotype, enabling comparisons despite the heteroscedasticity caused by disrupted canalization in mutant and transgenic lines. This decision was supported by a likelihood ratio test comparing models with uniform and genotype-specific φ values, as well as convergence checks (glmmTMB) and residual model fit (DHARMa 0.5.0) (*105*). Estimated marginal means and Šidák-adjusted p-values were found using emmeans 2.0.3 (*100*). All analysis scripts and processed data are available on GitHub (https://github.com/thecapitulab/AsterCLV3-qPCR-and-carpel-models) and archived at Zenodo (https://doi.org/10.5281/zenodo.22184594). Analyses were performed in R 4.6.0.

Lettuce (PI 251246) forms a paniculate capitulescence with a terminal capitulum that reaches anthesis first, followed by capitula terminating the secondary branches in basipetal order. To avoid variation due to developmental age, only capitula terminating secondary branches were analyzed in petal, stigma, and total floret counts for lettuce mutant and complementation experiments. Two capitula were quantified per plant in the initial WT vs *Lsclv3-cr* analyses, then four of the first six capitula were quantified in mutant complementation analyses. Stem width was measured at the time of floral organ counts using digital calipers around stem above the second true leaf.

### Protein expression and purification

The codon-optimized ectodomains of AtBAM1, AtBAM1^G199A^, AtBAM1^G199Q^ (AT5G65700, residues 20–629), AtBAM2 (AT3G49670, residues 23–623), LsBAM1/2a (LsBAM12a_g14072 (*26*), residues 19–620), HaCLV1b (HaCLV1b_Ha412HOChr14g00038256-RA.p1.prot, residues 76–682,), and LsCLV1a (g5782 (*26*), GenBank: PLY83950.1, residues 23–631) were cloned into a modified pFastBac vector (Geneva Biotech) under a polyhedrin promoter and fused to a Drosophila BIP signal peptide (*106*). The ectodomains were followed by a TEV (tobacco etch virus) protease cleavage site and a tandem affinity Strep-9xHis tag at their carboxy termini. Baculovirus generation was performed in DH10 cells, and virus amplification was done in *Spodoptera frugiperda* Sf9 cells. Proteins were produced in *Trichoplusia ni* Tnao38 cells infected at a multiplicity of infection (MOI) of 3. Cells were grown for 24 h at 28 °C followed by 48 h at 22 °C at 110 rpm. Secreted proteins were purified sequentially by Ni^2+^ affinity chromatography using HisTrap excel columns (Cytiva) equilibrated in 25 mM KPi, 500 mM NaCl, pH 7.8, followed by Strep-tag affinity chromatography using Strep-Tactin Superflow high-capacity columns (IBA, Germany) equilibrated in 25 mM Tris, 250 mM NaCl, 1 mM EDTA, pH 8.0. Affinity tags were removed by incubating the purified proteins with recombinant His-tagged TEV protease in a 1:50 (w/w) ratio overnight at 4 °C. Proteins were further purified, and cleaved tags were removed, by size-exclusion chromatography on a HiLoad 16/600 Superdex 200 column (Cytiva, 28-9893-35) equilibrated in 20 mM citric acid, 150 mM NaCl, pH 5.0. Peak fractions containing the target proteins were concentrated using Amicon Ultra 30 kDa concentrators (UFC9030). The wild-type BAM1 ectodomain was analyzed for purity and integrity by analytical SEC (size exclusion chromatography) with and SDS-PAGE (*43*). AtBAM1 pocket variants were analysed for purity and integrity by analytical SEC using a Bio-Rad ENrich SEC 650 column (7801650) and SDS-PAGE (Supplemental Fig. 12c). AtBAM2 and LsBAM1/2a were analysed by analytical SEC using a Superdex 200 10/300 GL column (Cytiva) and SDS-PAGE (Supplemental Fig. 13b). HaCLV1b and LsCLV1a were analysed by analytical SEC using a Superdex 200 10/300 GL column (Cytiva) and SDS-PAGE (Supplemental Fig. 18, d and e, respectively).

### Isothermal titration calorimetry (ITC)

Binding experiments were performed at 25 °C using a MicroCal PEAQ-ITC (Malvern Instruments) with a 200 µL standard cell and a 40 μL titration syringe. Protein ectodomains were gel filtrated into the ITC buffer (20 mM sodium citrate, 150 mM NaCl, pH 5.0). Protein concentrations were determined based on their molar extinction coefficients and calculated molecular weights after removal of the signal peptide and affinity tags. A typical experiment consisted of injecting 2 μL or 3 μL of a range between 100 and 1200 μM solution of CLE peptides and variants, into a 10 to 30 μM receptor protein in solution in the cell. Experiments were run at 150 s intervals and 500 rpm stirring speed. Experiments were done in duplicates and data were analyzed using the MicroCal PEAQ-ITC Analysis Software provided by the manufacturer. Control experiments of ligand vs buffer were performed independently, and no dilution heat was observed. All ITC runs used for data analysis have an N ranging from 0.9 to 1.1. The N values were fitted to 1 in the analysis. Control experiments of ligand vs buffer were performed independently, and no dilution heat was observed. All ITC runs used for data analysis have an N ranging from 0.9 to 1.1. The N values were fitted to 1 in the analysis.

### Structural modeling and prediction of water molecules within the binding pocket

Structural models of BAM1/CLV1 receptors in complex with peptides were generated using the Alphafold3 server (https://alphafoldserver.com/) (*107*). For AtCLE13p in complex with AtBAM1, final peptide-receptor refinement was performed using high-resolution modeling through the Rosetta FlexPepDock tool (*108*), with total energy score of -430.181 and a RMSD of 0.997. The prediction of water molecules within the binding pocket was done using Waterdock 2.0 (*109*). Generation of molecular images was performed using ChimeraX (*110*) and PyMOL (The PyMOL Molecular Graphics System, Version 2.6.1 Schrödinger, LLC.).

### Conservation analysis of receptors surface

Conservation analysis within BAMs, CLVs, and TDRs receptors was made using ESPript 3.0 server (https://espript.ibcp.fr/ESPript/ESPript/) (*111*). A multiple sequence alignment produced with Clustal Omega(*112*) was used as input. Images were generated using PyMOL (The PyMOL Molecular Graphics System, Version 3.0 Schrödinger, LLC).

## Supplementary Figures

**Supplementary Fig. 1:**
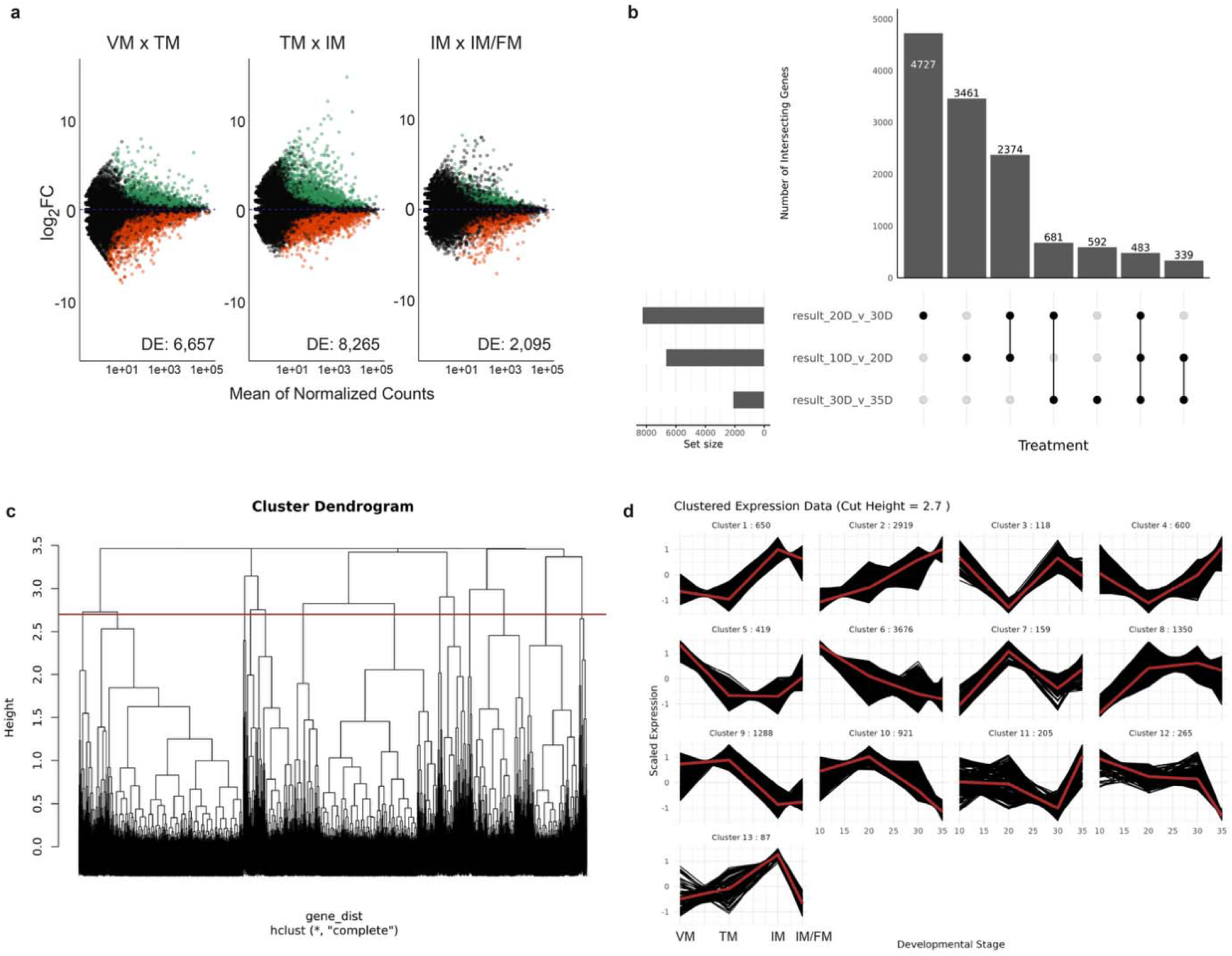
Transcriptomic analysis of sunflower shoot meristems throughout capitulum development. a, Volcano plots showing differentially expressed (DE) genes across pairwise comparisons of sunflower inflorescence development (adj. p-value <0.05), total number of DE genes indicated for each comparison (green = up; red = down). b, Upset plot of differentially expressed (DE) genes across pairwise comparisons showing shared and unique DE genes. c, Dendrogram of hierarchical clustering. Red line at height 2.7 indicates the cut site used in downstream clustering analyses. d, Co-expressed gene clusters identified at the 2.7 cut site. Plots show the scaled expression of each gene in the cluster with the red line representing the average for the cluster. Number of genes in each cluster indicated with each graph. Lists of genes in each cluster are presented in Supplementary File 1.

**Supplementary Fig. 2:**
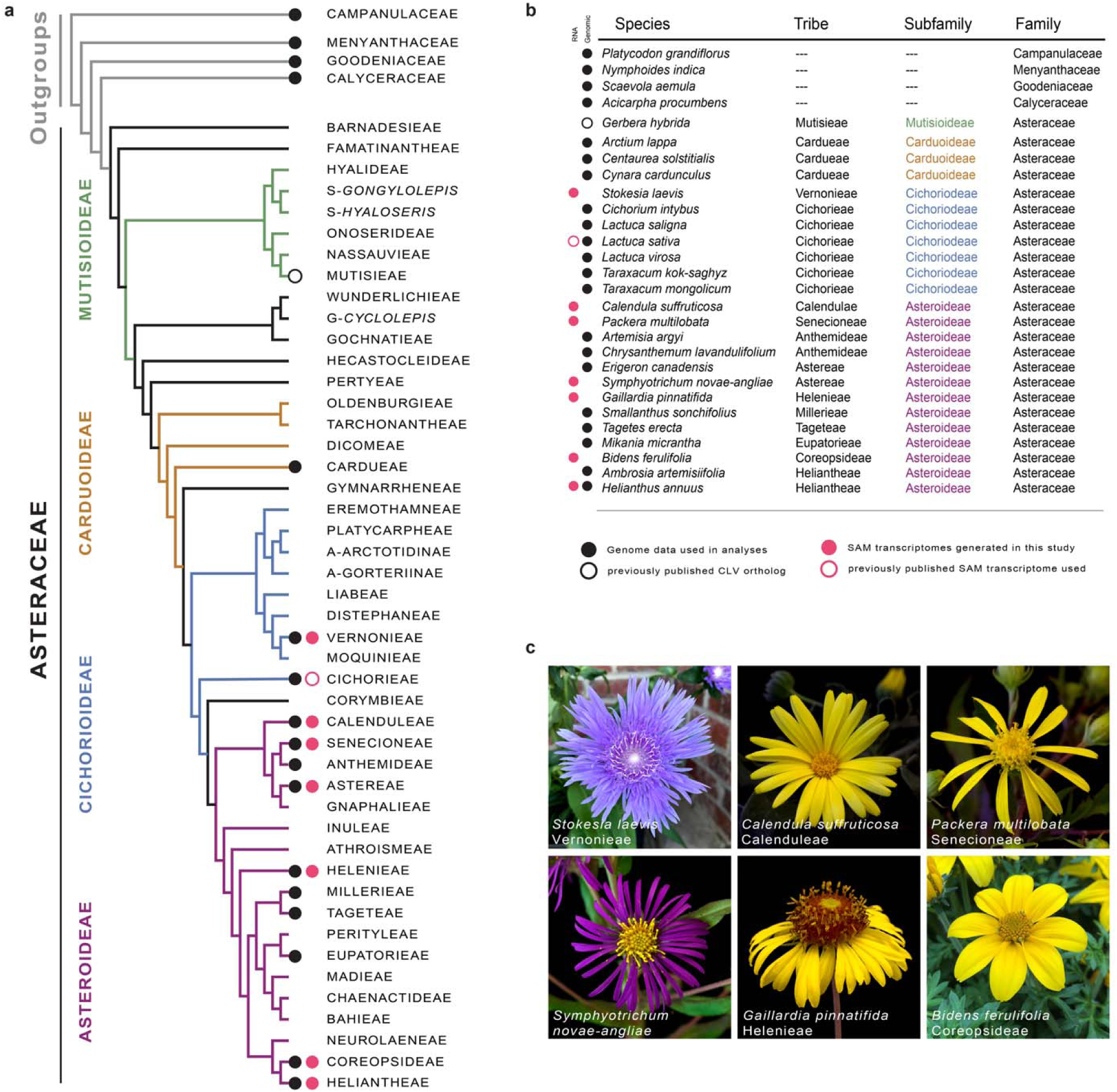
Taxonomic sampling of species across Asterales for gene identification. **a**-b, A family/tribe-level phylogenic tree and table showing the Asteraceae and immediate outgroups along with species used in this study. Four subfamilies in the Asteraceae are highlighted: Mutisioideae (green), Carduoideae (brown), Cichorioideae (blue) and the Asteroideae (purple). Type of data used in our analyses is indicated with black (genomic) or red (SAM transcriptomic) circles. c, Representative pictures of capitula from the six species for which *de novo* transcriptomes were assembled from SAMs spanning floral development (see methods).

**Supplementary Fig. 3:**
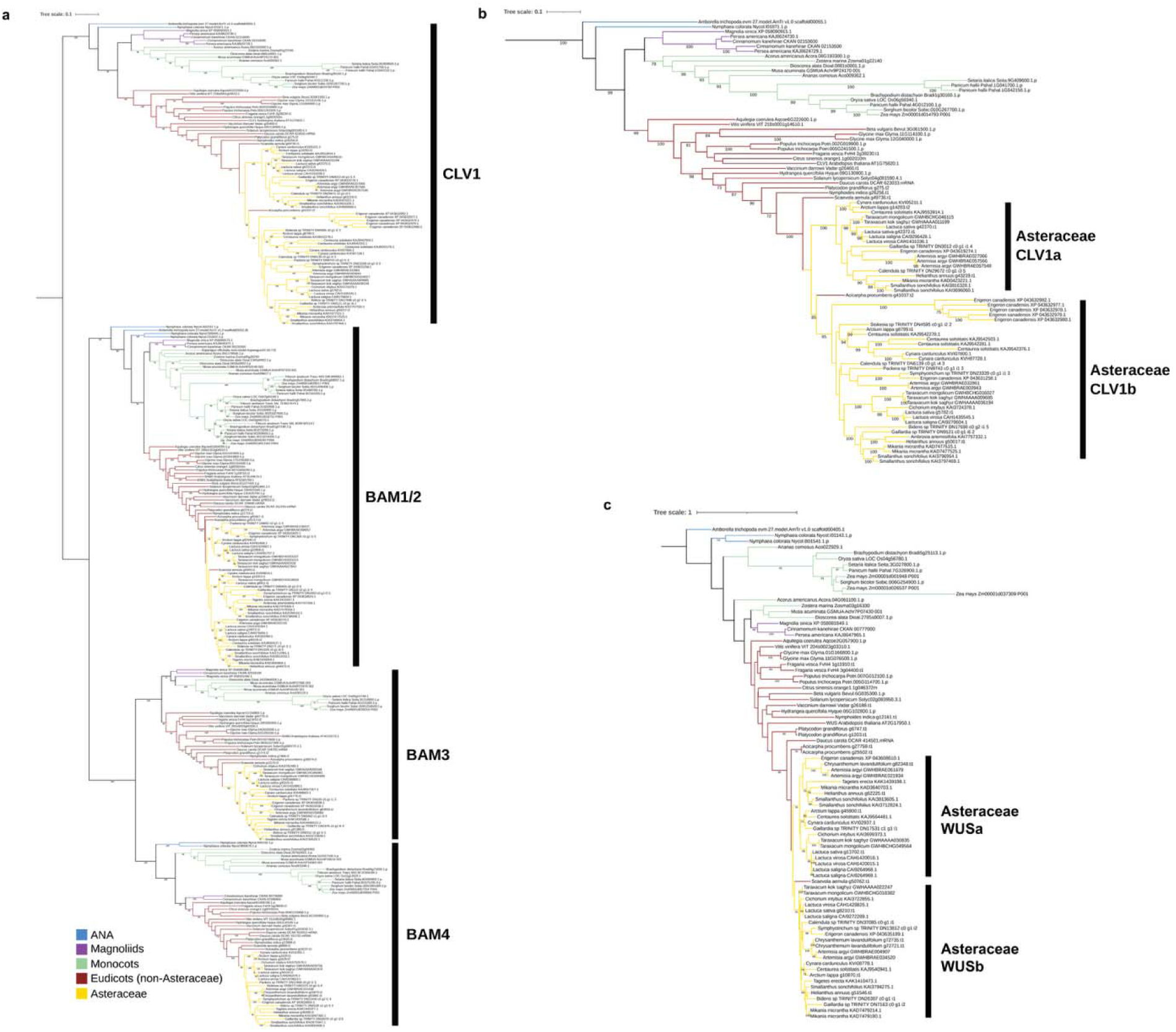
CLV1/BAM receptors and WUS gene trees. a, RAxML phylogenetic tree of the CLAVATA1/BARELY ANY MERSITEM (CLV1/BAM) related receptors, with clades labeled with known *A. thaliana* and *Solanum lycopersicum* (tomato) orthologs. b, expanded view of CLV1 orthologs across flowering plants showing an Asteraceae specific duplication (CLV1a and CLV1b clades). c, WUSCHEL (WUS) orthologs across flowering plants revealing an Asteraceae specific duplication (WUSa and WUSb clades). *WUSb* orthologs were all expressed in sunflower, lettuce and Gerbera shoot meristems and was identified in 4/6 SAM transcriptomes assembled for this study (Fig. 1f; Supplementary Fig. 14b). *HaWUSb* expression was also downregulated in response to CLV3p treatments (Fig 5k). a-c, Trees were all built from protein alignments and branch support was estimated with 200 rapid bootstrap replicates (see methods). Branches colored based on taxonomy. High resolution version of this figure can be found in Supplementary Files.

**Supplementary Fig. 4:**
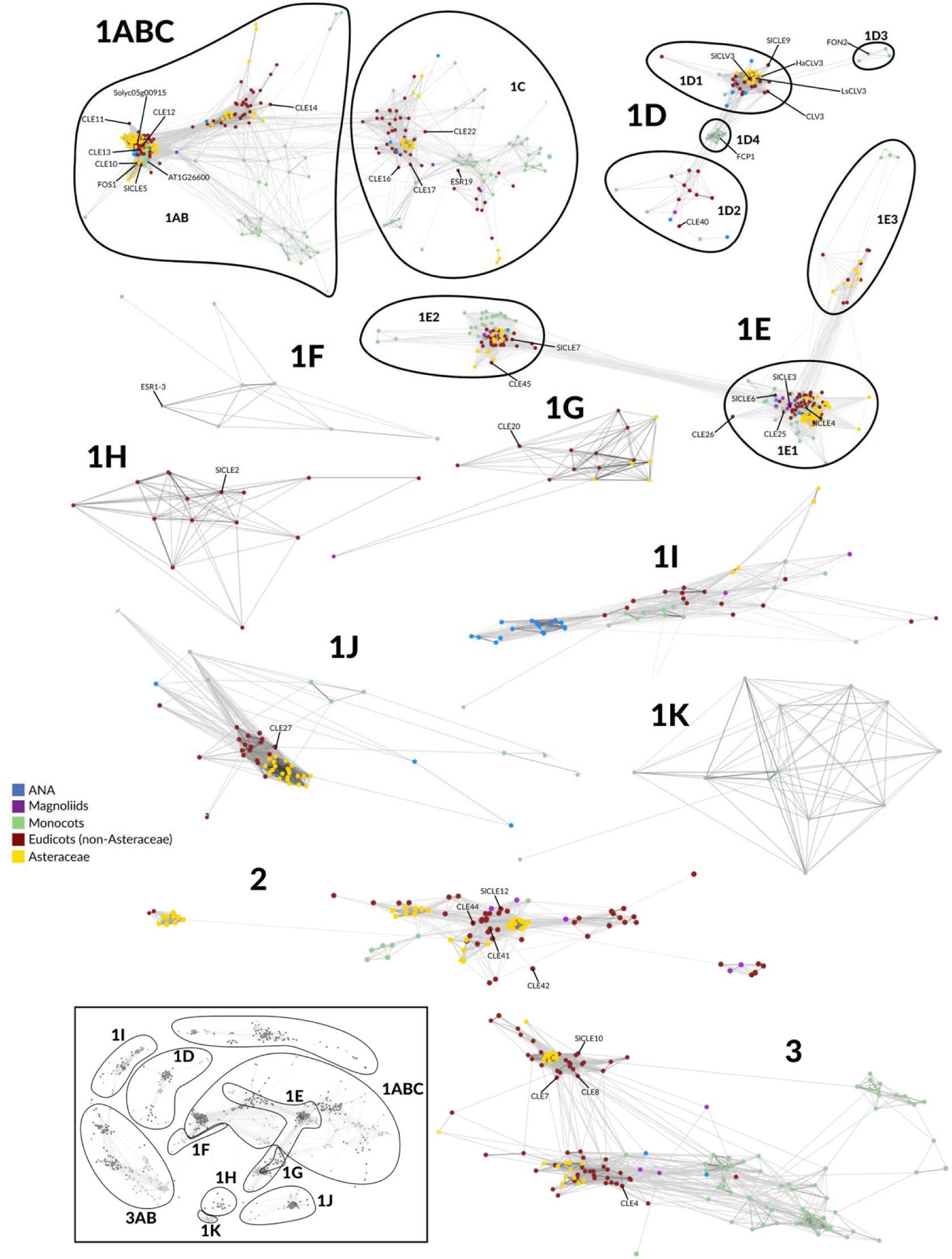
CLE clustering and identification. a, CLANs-based clustering of CLV3/EMBRYO SURROUNDING REGION (CLE) pre-propeptides across flowering plants. Clusters were identified and extracted using a previously published CLEp-classification system (Goad *et al.* 2017). Genes colored based on taxonomy and known *A. thaliana* and *Solanum lycopersicum* (tomato) orthologs from each cluster are indicated (“CLE” for *A. thaliana* and “SlCLE” for tomato). Related to Supplementary File 3.

**Supplementary Fig. 5:**
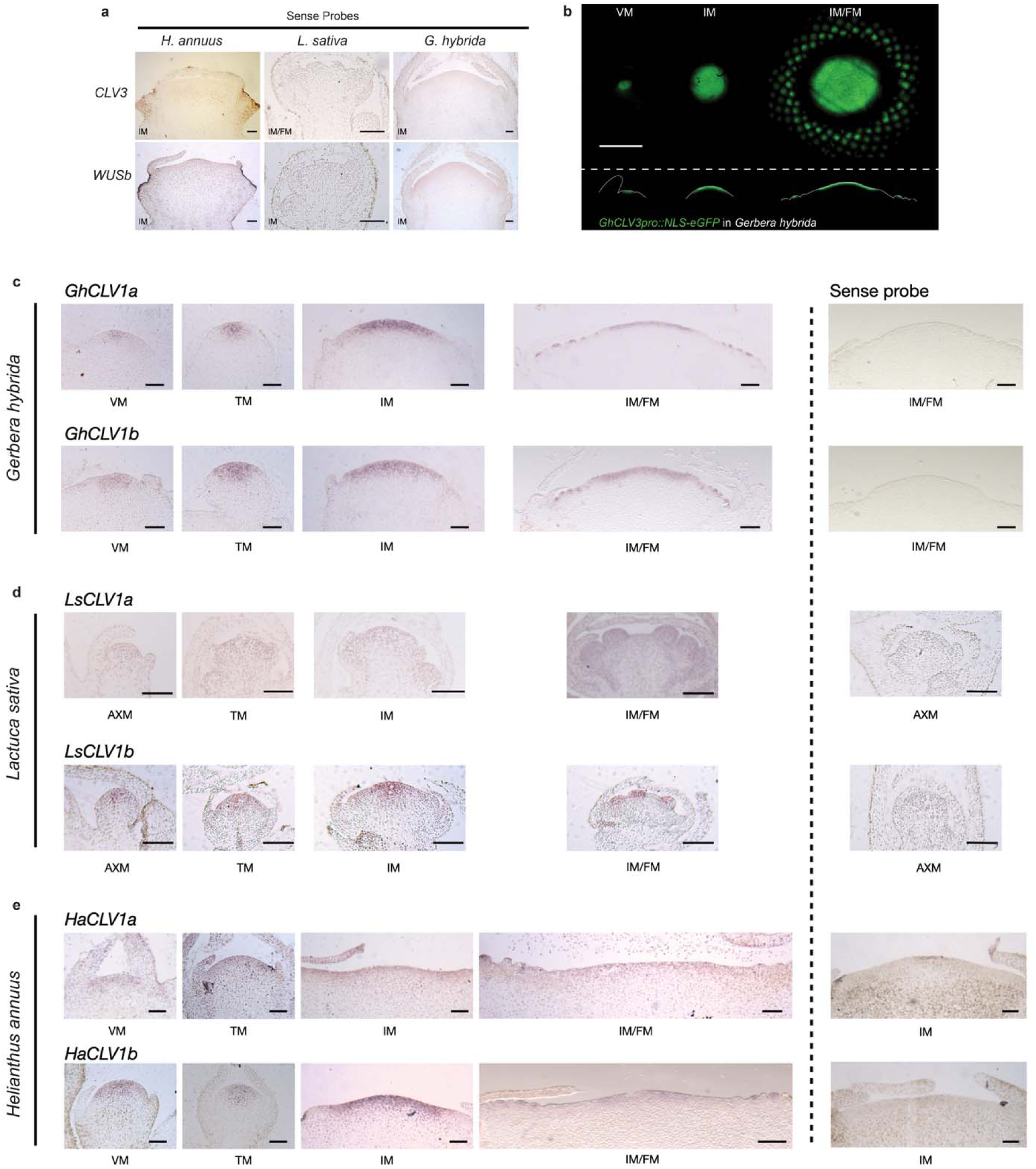
*CLAVATA* pathway *in situ* hybridization in developing Gerbera, lettuce and sunflower shoot meristems. a, sense probe hybridizations for *CLV3* and *WUSb* (related to Fig. 1f). b, Confocal micrographs of the *GhCLV3pro::NLS-eGFP* reporter in shoot meristems of *Gerbera hybrida* across inflorescence development. Upper panel shows a top-down view (xy-axis) and lower panel shows representative longitudinal sections (z-axis) of each shoot meristem with meristems outlined in white. c-e, analysis of *CLV1a* and *CLV1b* expression in Gerbera, lettuce and sunflower shoot meristems spanning vegetative (SAM) to floral development (IM and FM). Antisense probes on left and sense probe controls shown on right. Scale bars - 100µm (a, c-e) and 500µm (b).

**Supplementary Fig. 6:**
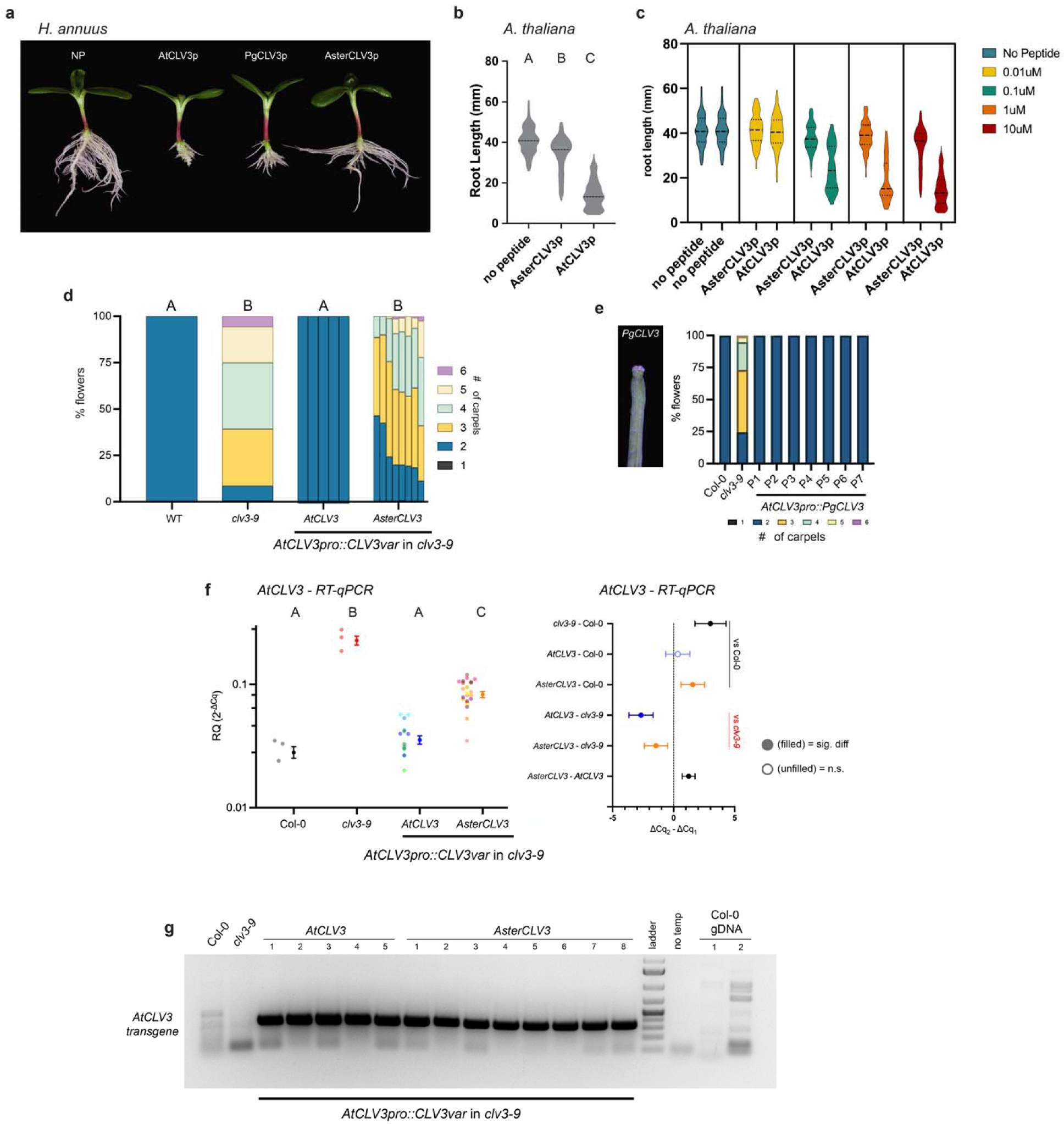
Functional analysis of AsterCLV3 in sunflower and *A. thaliana*. a, CLE peptide root assays showing *Helianthus annuus* (sunflower) response to 0.1μM AtCLV3p, PgCLV3p (*P. grandiflorus*), and AsterCLV3p compared to no peptide (NP) control. b-c, CLE peptide root assays showing *A. thaliana* (Col-0) response to a concentration gradient of AtCLV3p and AsterCLV3p compared to NP control. b, extracted 10μM peptide treatment for statistical comparison. Statistical summaries from one-way ANOVA and Tukey’s multiple comparison tests where A-C groups are significantly different (adj. p-value <0.0001). d, data from T3 individuals in complementation analysis of carpel number in *clv3-9* using a native *AtCLV3* promoter::gene (*AtCLV3pro::AtCLV3*) constructs with AtCLV3 and AsterCLV3 CLE domains swapped in (related Fig. 2h-i). Statistical summaries from Conway-Maxwell (COM)-Poisson GLMM with A-B groups significantly different (p ≤ 1.12E-07). e, T1 complementation analysis of carpel number in *clv3-9* of *AtCLV3pro::PgCLV3* (across 7 independent transformation events). f, RT-qPCR analysis of *AtCLV3* expression in Col-0, *clv3-9*, *AtCLV3pro::AtCLV3* and *AtCLV3pro::AsterCLV3* T3 homozygous lines. Left shows plot with mean and SEM (RQ 2^-ΔCq^) expression with all biological replicates shown as individual dots. Statistical summaries from restricted maximum likelihood linear mixed model with A-C groups significantly different (p ≤ 0.0049). Right shows a forest plot of pairwise comparisons with 95% CI of mean difference (ΔCq_2_- ΔCq_1_); effect sizes reported as Hedge’s g in Supplementary File 8. g, agarose gel electrophoresis showing primers amplifying *AtCLV3pro::CLV3var* transgene from cDNA of multiple independent lines used in d. Col-0 cDNA, *clv3-9* cDNA, Col-0 gDNA and no template reactions used as negative controls. b-c (n=individual roots analyzed), in b-c, n= 132 (NP), 139 (0.01µM AtCLV3p), 131 (0.1µM AtCLV3p), 142 (1µM AtCLV3p), 139 (10µM AtCLV3p), 128 (0.01µM AsterCLV3p), 123 (0.1µM AsterCLV3p), 121 (1µM AsterCLV3p), 125 (10µM AsterCLV3p). In d, (n=individual plants with 18-20 flowers counted on each) n= 6-8 plants per line. In e, n=individual plants with 10 flowers counted on each, 7 independent lines. In f, n=3-4 biological replicates of 100mg of seedling tissues for each transgenic line. See complete statistical analyses and sample sizes in Supplementary File 8.

**Supplementary Fig. 7:**
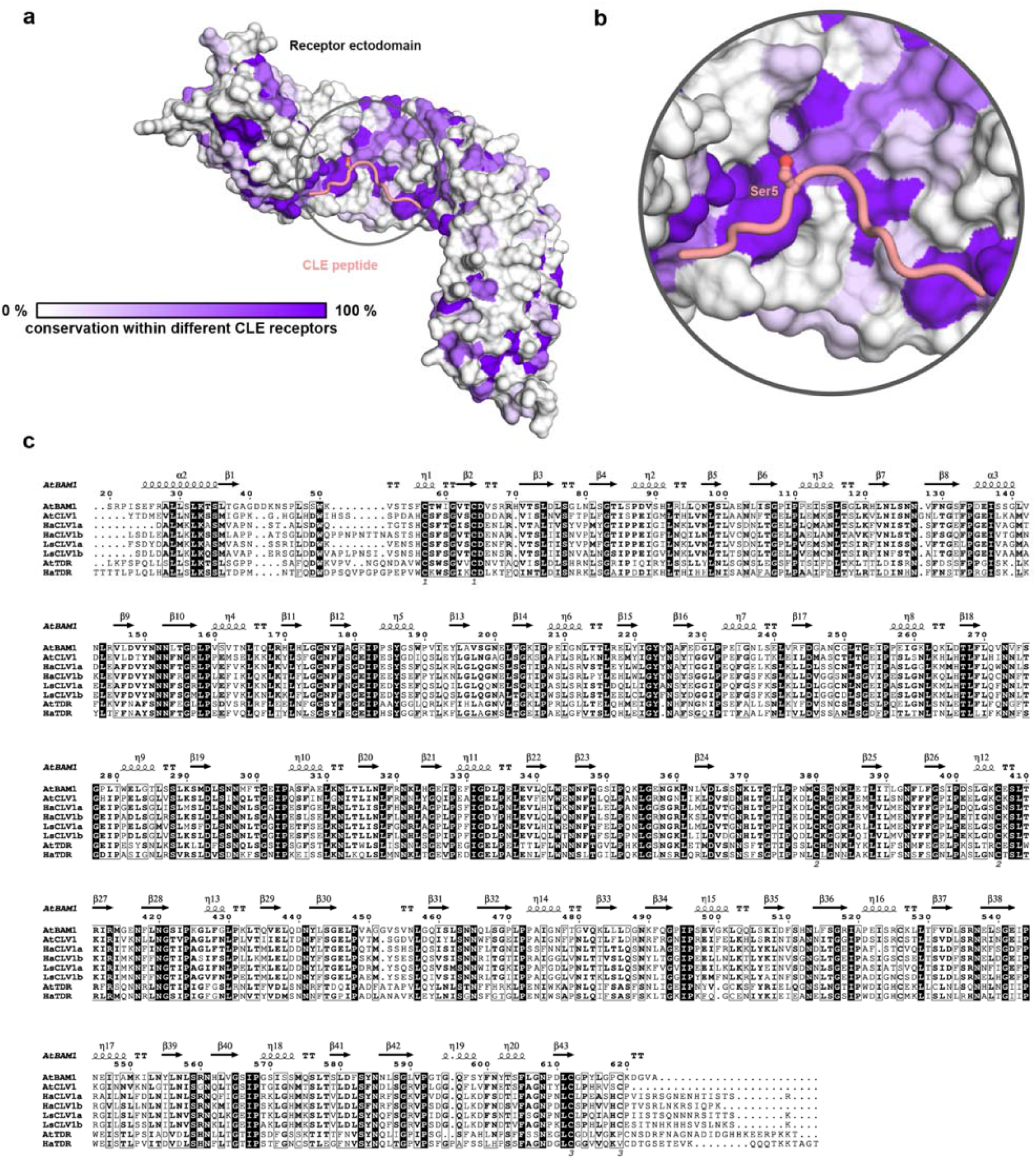
Structural comparison between TDR/PXY and BAM1/CLV receptors reveals lower conservation in their peptide-binding pockets. a, Surface representation of BAM1/CLV and TDR/PXY receptor ectodomains, colored based on sequence conservation. b, Close-up view of the peptide-binding pocket, highlighting the differences in conserved residues. c, Sequence alignment of the receptors analyzed in (a), illustrating conservation patterns across species.

**Supplementary Fig. 8:**
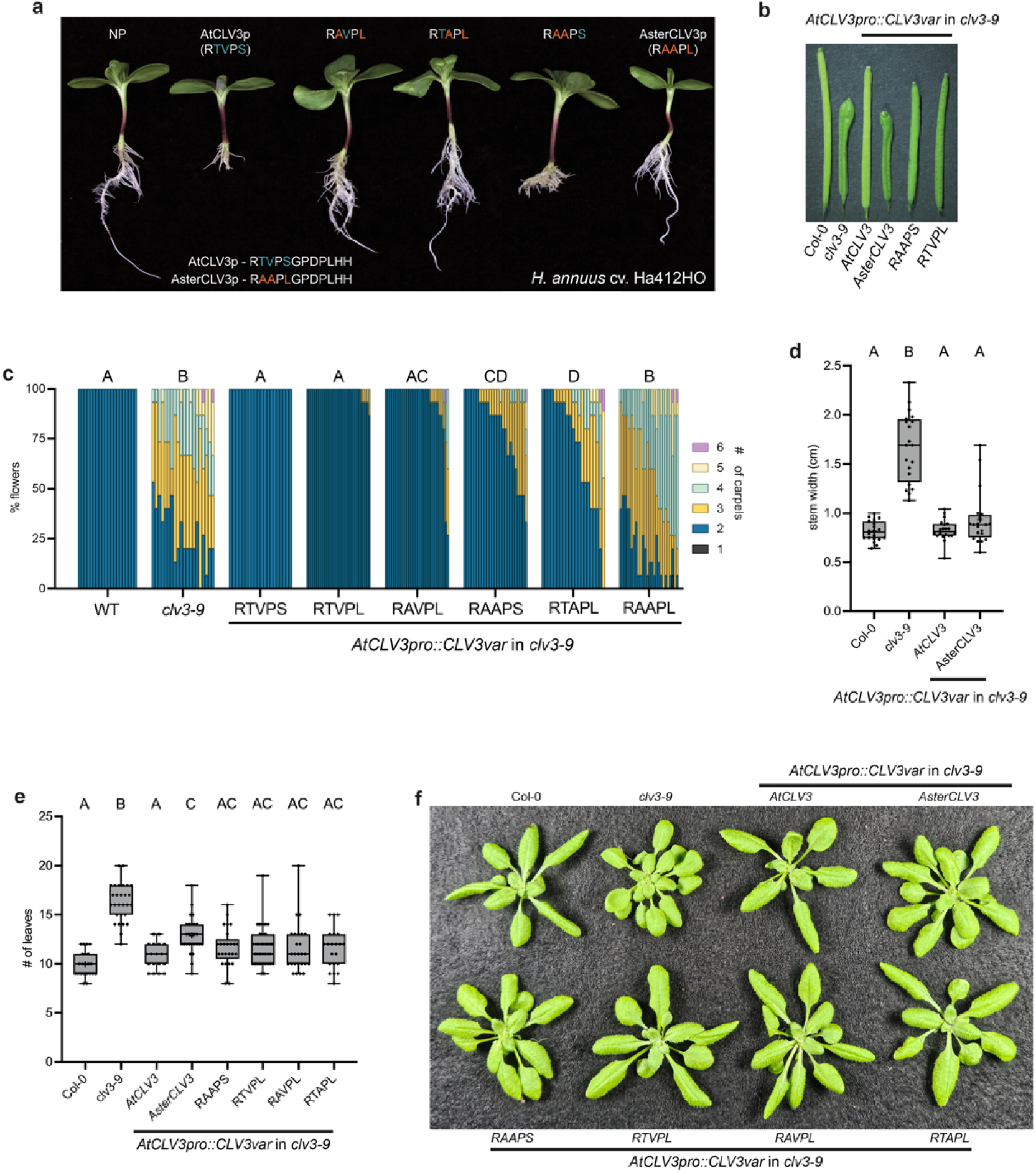
Functional analysis of CLV3 variants in sunflower and *A. thaliana*. a, CLE peptide root elongation assays showing responses to 0.1µM AtCLV3p, AsterCLV3p and intermediate CLV3p variants in sunflower. b-c, T1 complementation analysis of carpel number in *clv3-9* using a native *AtCLV3* promoter::gene (*AtCLV3pro::AtCLV3*) constructs with AtCLV3, RTVPL, RAVPL, RAAPS, RTAPL and AsterCLV3 CLE domains swapped in. b, representative *A. thaliana* siliques from *AtCLV3pro::CLV3var* in *clv3-9* complementation. c, summary of carpel number percentages across CLV3var comp lines. Statistical summaries from Conway-Maxwell (COM)-Poisson GLMM with A-D groups significantly different (p ≤ 0.0017). d, stem width from T1 complementation *AtCLV3pro::AtCLV3* and *AsterCLV3* in *clv3-9.* Statistical summary from one-way ANOVA where A and B groups are significantly different (adj. p-value < 0.0001). e-f, rosette leaf counts from *AtCLV3pro::CLV3var* in *clv3-9* complementation. Statistical summaries from Kruskal-Wallis and Dunn’s multiple comparison tests where A-C groups are significantly different (adj. p-value <0.05). In c, (n=individual plants with 15 flowers counted each) n= 20 (Col-0), n= 20 (*clv3-9*), 20 (*AtCLV3*), 33 (*RTVPL*), 30 (*RAVPL*), 25 (*RAAPS*), 21 (*RTAPL*), and 26 (*AsterCLV3*). In d, (n=stem measurements from individual plants), n= 20 for each. In e, (n=rosette leaf number from individual plants) n= 22 (Col-0), 28 (*clv3-9*), 19 (*AtCLV3*), 27 (*AsterCLV3*), 25 (*RAAPS*), 35 (*RTVPL*), 26 (*RAVPL*), and 21 (*RTAPL*).

**Supplementary Fig. 9:**
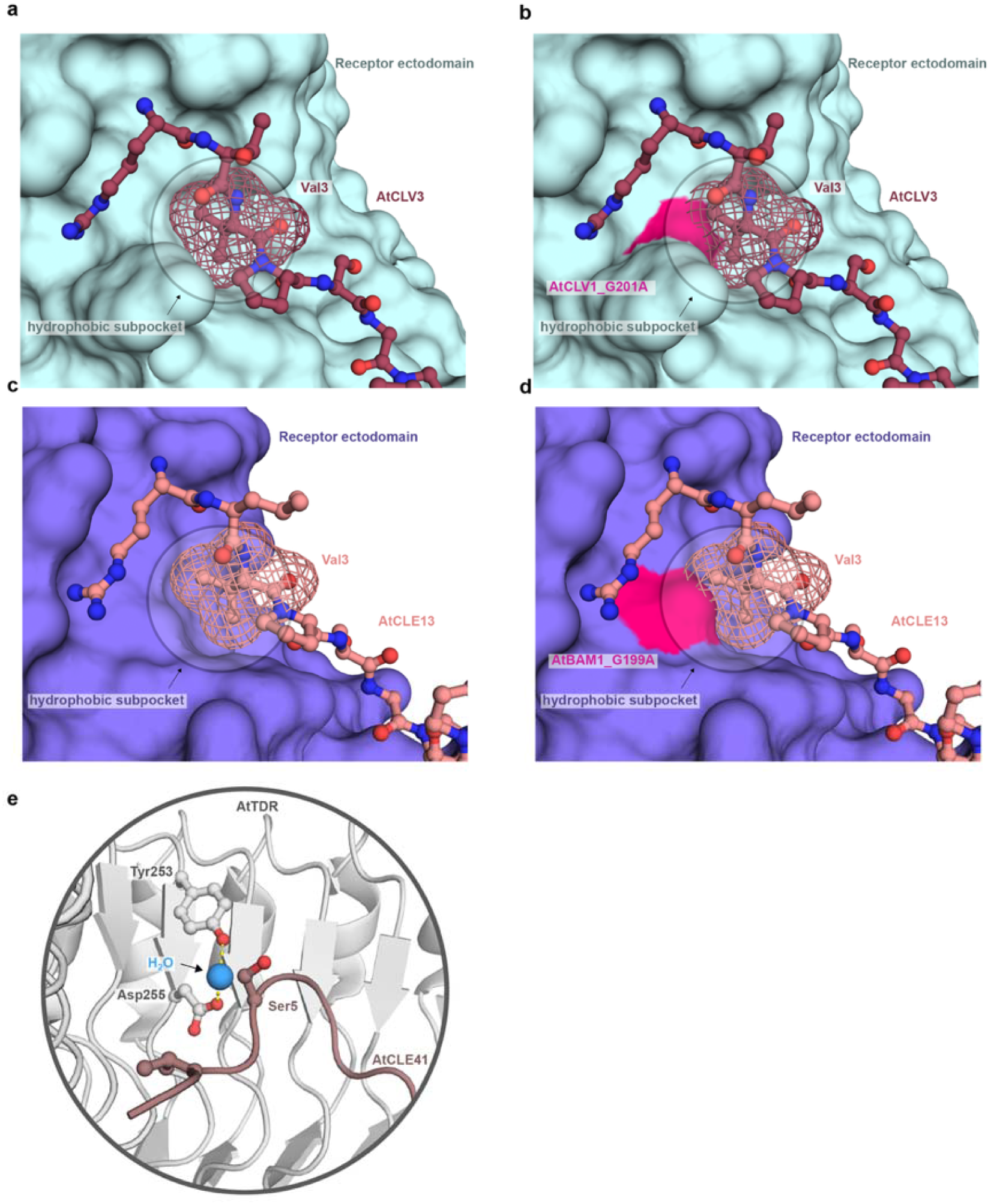
BAM/CLV receptors share a conserved hydrophobic subpocket that anchors the peptide within the binding groove. a-d, Structural representation of the N-terminal peptide-binding site in the AtCLV1-AtCLV3p (a-b) and AtBAM1-AtCLE13p (c-d) complex. The side chain of AtCLV3p/AtCLE13p Val3 fits into both receptors’ hydrophobic subpockets. The mesh represents the volume occupied by Val3 atoms. b and d, Structural model of AtCLV1 G201A and AtBAM1 G199A variants, highlighting the hydrophobic cavity. Ala substitutions reduce the available space for AtCLV3p/AtCLE13p Val3, causing steric clashes. AtCLV1 (light blue) and AtBAM1 (purple) are shown in surface representation, while AtCLV3p (dark red) and AtCLE13p (pink) are depicted in stick-and-ball representation. e, Close-up view of the binding pocket of AtTDR/PXY receptor in complex with AtCLE41p (PDB ID: 5GR9). Despite structural differences between BAM/CLV and TDR/PXY receptors, a water molecule is positioned near AtCLE41p Ser5, coordinating water-mediated interactions between the conserved Tyr253 and Asp255 and AtCLE41p peptide backbone.

**Supplementary Fig. 10:**
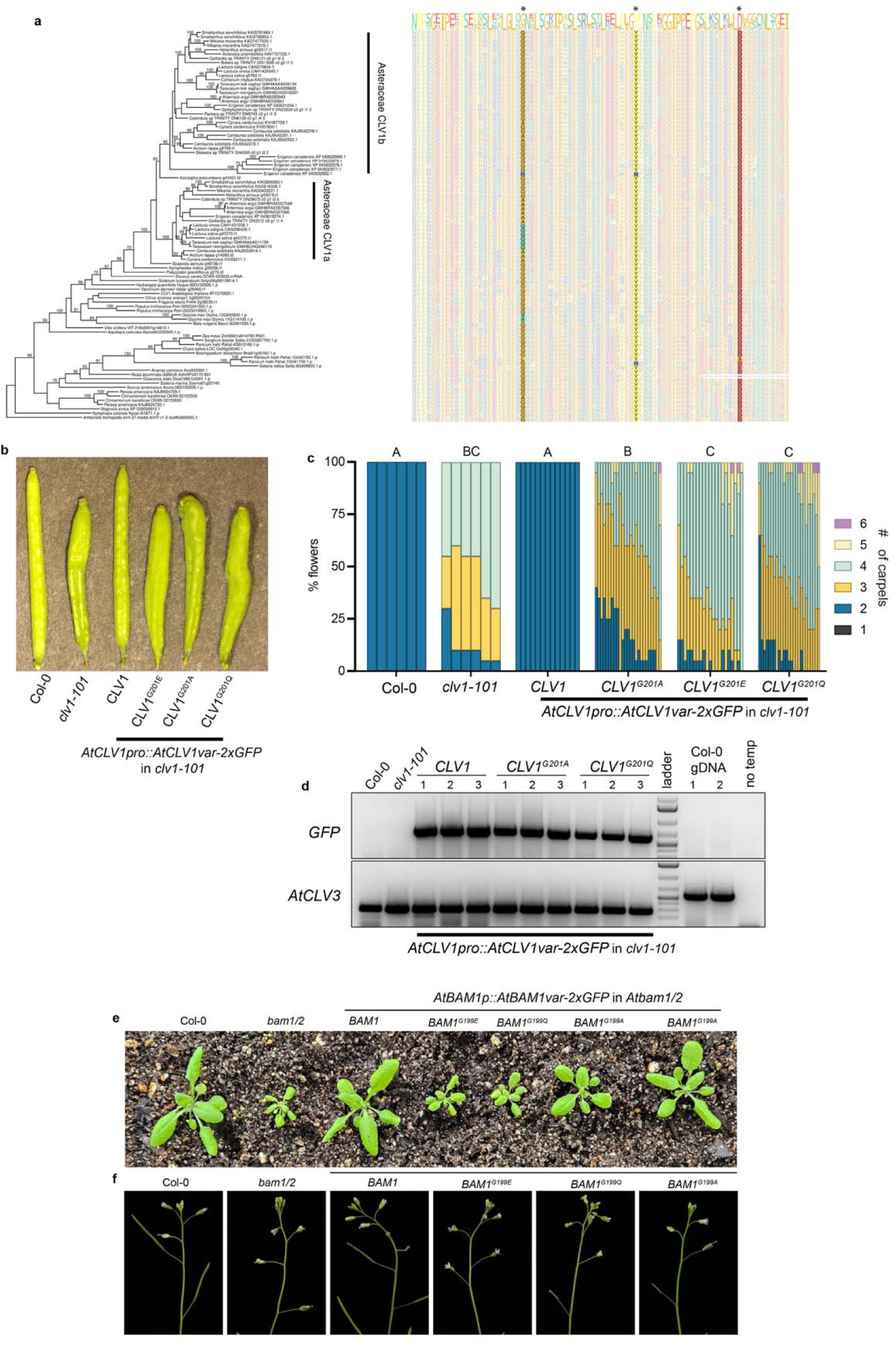
Functional analysis of AtBAM1/AtCLV1’s peptide-binding pocket. a, Sequence alignment of the conserved CLV1 peptide binding domain with three critical residues highlighted (G201, Y225, and D247 – numbering based on AtCLV1 position). b, representative images of *A. thaliana* siliques and breakdown data of individuals from T1 complementation analysis of carpel number in *clv1-101* using a native *AtCLV1* promoter::gene (*AtCLV1pro::AtCLV1*) constructs with AtCLV1, AtCLV1^G201Q^, AtCLV1^G201E^ and AtCLV1^G201A^. c, T1 complementation analysis of carpel number in *clv1-101* using a native *AtCLV1* promoter::gene (*AtCLV1pro::AtCLV1*) construct expressing AtCLV1, AtCLV1^G201Q^, AtCLV1^G201E^ and AtCLV1^G201A^. Statistical summaries from Conway-Maxwell (COM)-Poisson model with A-C groups significantly different (p ≤ 4.96E-06). d, agarose gel electrophoresis showing primers amplifying C-terminal GFP fusion on *AtCLV1pro::AtCLV1* constructs (AtCLV1, AtCLV1^G201Q^ and AtCLV1^G201A^) from cDNA of multiple independent lines used in c. Col-0 cDNA, *clv1-101* cDNA, Col-0 gDNA and no template reactions used as negative controls, and *AtCLV3* amplification used as a positive control of shoot meristem derived cDNA across all samples. e, Representative images of *A. thaliana bam1/2* rosette size phenotype T1 complementation using *AtBAM1* promoter::gene (*AtBAM1::AtBAM1*) constructs with AtBAM1, AtBAM1^G199E^, AtBAM1^G199Q^ and AtBAM1^G199A^. f, Representative images of *A. thaliana bam1/2* sterility phenotype with AtBAM1, AtBAM1^G199E^, AtBAM1^G199Q^ and AtBAM1^G199A^ lines from e. In c, (n=plants with 20 flowers counted on each) n= 6 (Col-0), 6 (*clv1-101*), 13 (*AtCLV1*), 25 (*AtCLV1^G201A^*), 21 (*AtCLV1^G201E^*), 25 (*AtCLV1^G201Q^*). High resolution version of Supplementary Fig. 10a can be found in Supplementary Files.

**Supplementary Fig. 11:**
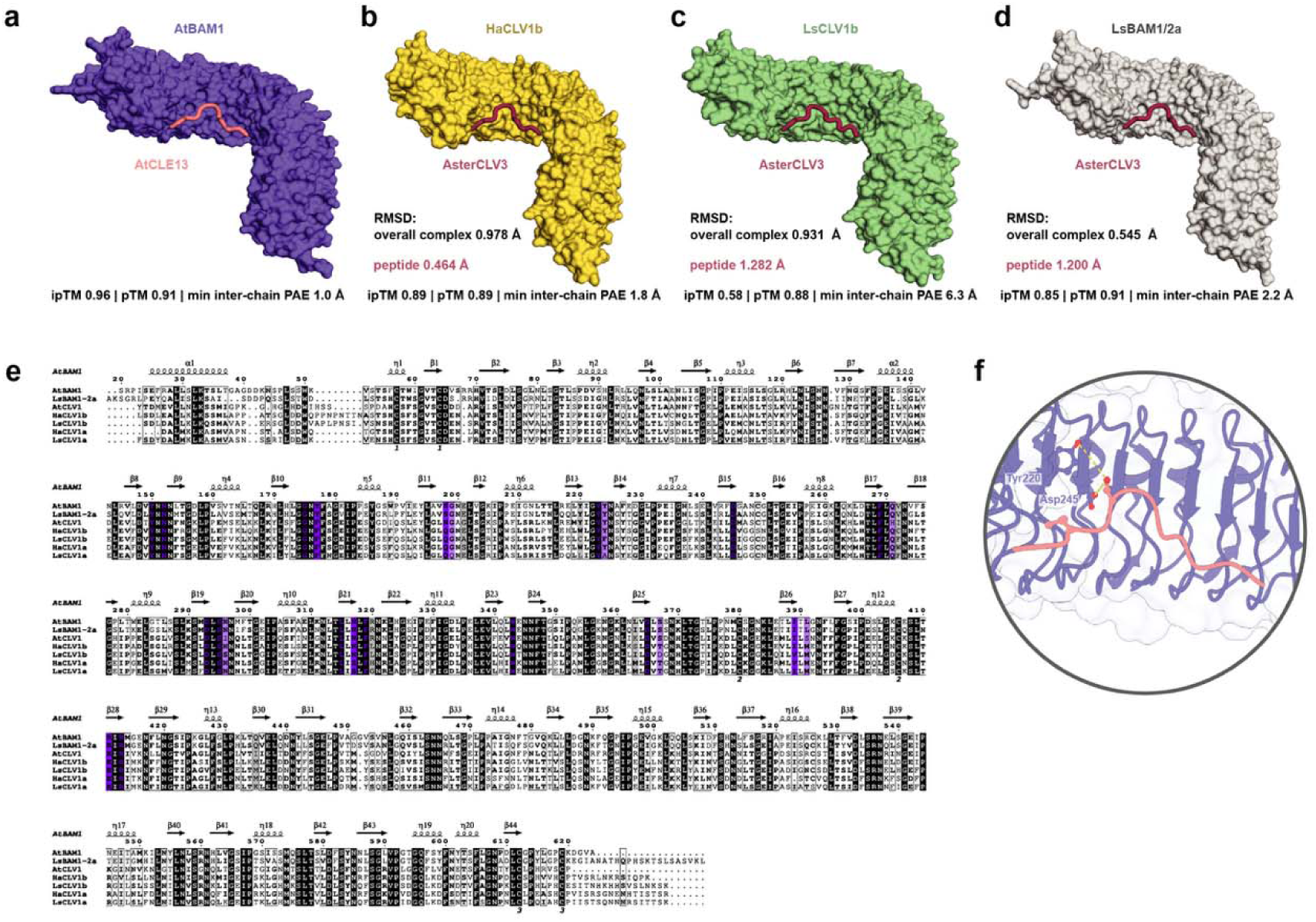
BAM1 and CLV1 share a conserved peptide-binding pocket. a-d, AlphaFold 3 structural models of *A. thaliana* BAM1 (AtBAM1), *Helianthus annus* CLV1b (HaCLV1b), *Lactuca sativa* CLV1b (LsCLV1b) and LsBAM1/2a in complex with CLV/CLE peptides. Overall and peptide RMSDs (root mean squared deviations) relative to AtBAM1-CLE13 are indicated. Model confidence is indicated by ipTM (interface predicted template modeling score), pTM (predicted template modeling score), and minimum inter-chain PAE (predicted aligned error, Å), according to AlphaFold 3 metrics. e, Sequence alignment of the ectodomains of BAM1 and CLV1 receptors across species analyzed in Fig. 4a. The residues forming the pocket and involved in peptide coordination are highlighted according to the conservation algorithm used by ESPript 3.0 server (https://espript.ibcp.fr/ESPript/ESPript/) (110) f, AlphaFold 3 model showing AtBAM1 residues Tyr220 and Asp245 (purple sticks) positioned near CLE13 Ser5 (pink sticks), create a local environment supporting additional polar interactions. Distances are indicated by yellow dashed lines within a distance of 5.8 Å. Close-up view of the receptor binding pocket (purple cartoon) and the CLE peptide in pink. High resolution version of this figure can be found in Supplementary Files.

**Supplementary Fig. 12:**
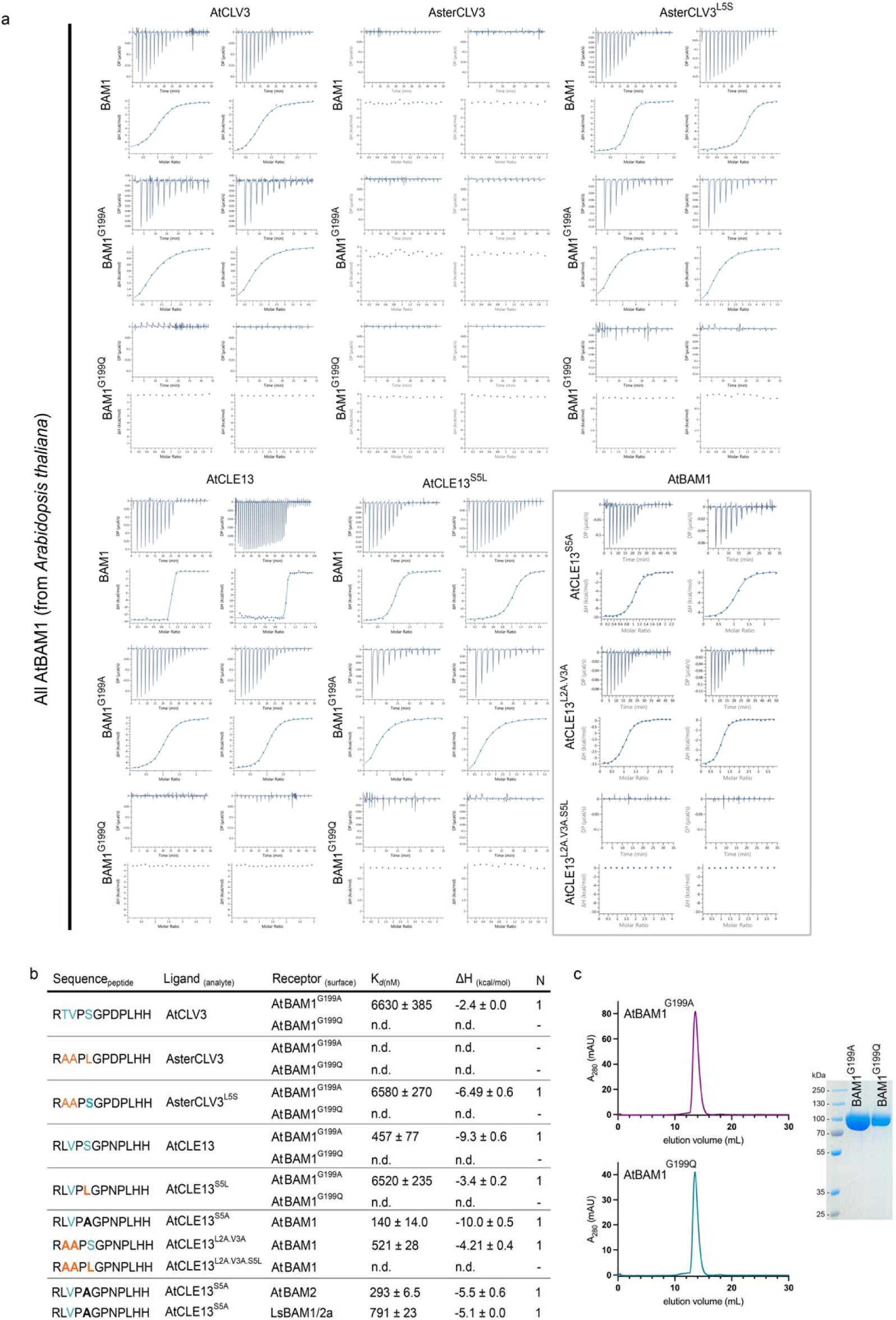
AtBAM1/peptide binding assays. Isothermal titration calorimetry (ITC) assays of AtBAM1 and its mutant variants binding to different CLV/CLE peptides. Shown are the ITC thermograms from independent experiments conducted for each peptide, corresponding to the analysis presented in Fig. 4. b, ITC summary table of CLE/CLV peptides binding to AtBAM1, AtBAM1 variants, AtBAM2 and LsBAM1/2a ectodomains. K*_d_* (nM) represents binding affinity, ΔH (kcal/mol) indicates reaction enthalpy, and N denotes stoichiometry (n = 1 for a 1:1 interaction). Data are mean ± SD from at least two independent experiments, with undetected binding shown as n.d. c, Protein integrity and purity of BAM1 pocket variants. SEC analysis and SDS-PAGE of BAM1^G199A^ and BAM1^G199Q^.

**Supplementary Fig. 13:**
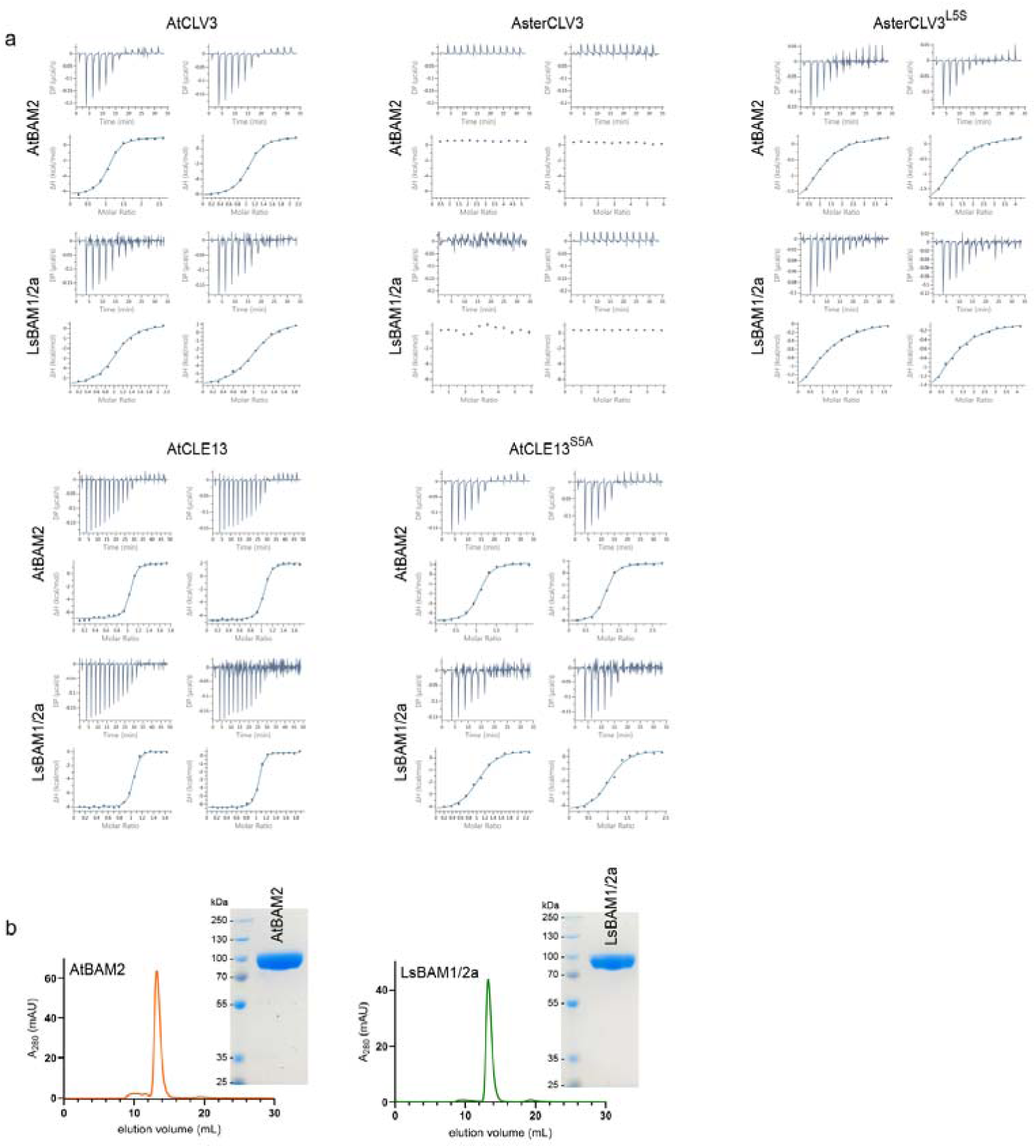
AtBAM2 and LsBAM1/2a peptide binding assays. Isothermal titration calorimetry (ITC) assays of AtBAM2 and LsBAM1/2a binding to different CLV/CLE peptides. Shown are the ITC thermograms from independent experiments conducted for each peptide. b, Protein integrity and purity. SEC analysis and SDS-PAGE of AtBAM2 and LsBAM1/2a ectodomains.

**Supplementary Fig. 14:**
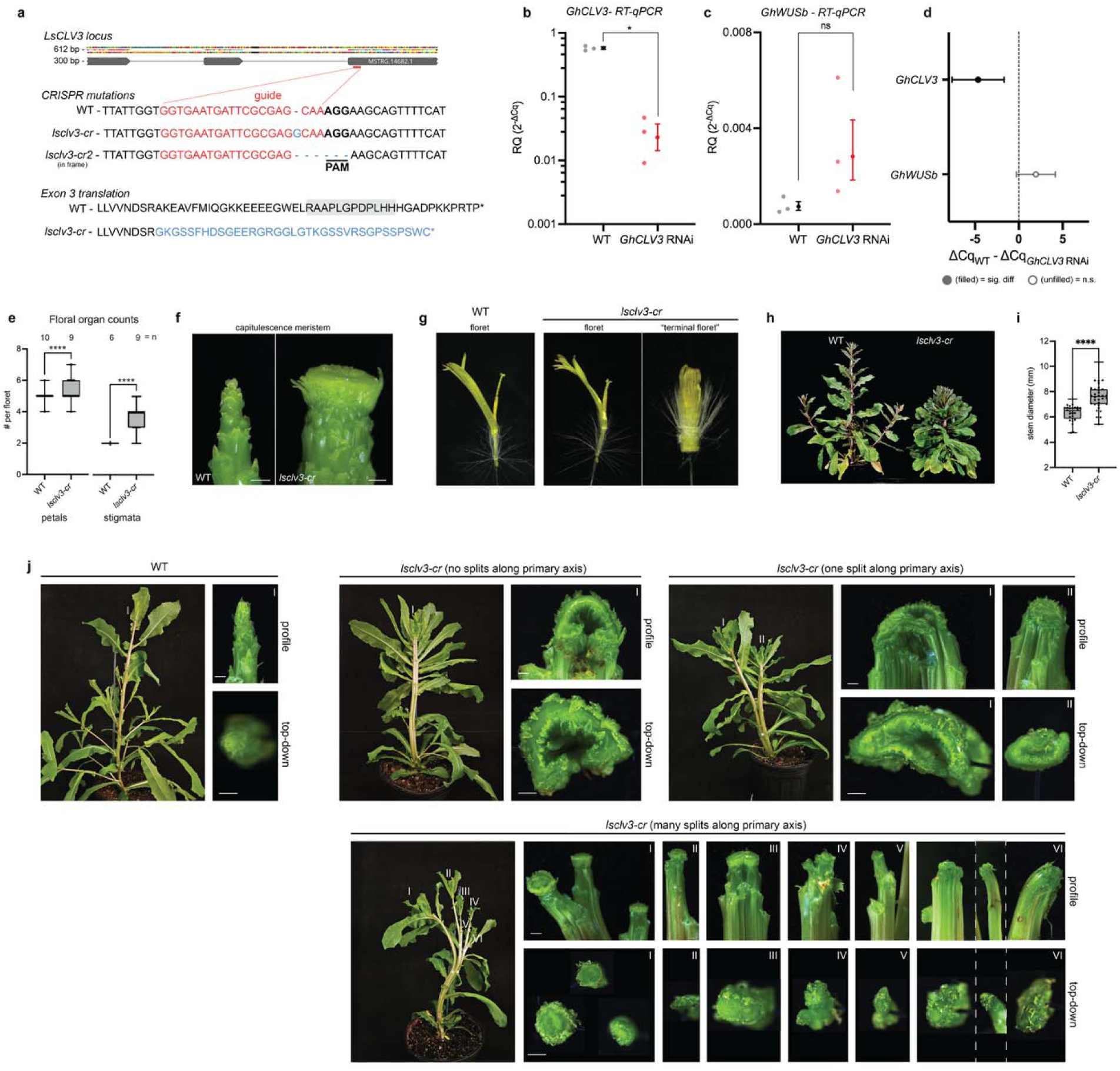
*CLV3* has conserved stem cell repressive function in lettuce and Gerbera. a, *LsCLV3* gene model with CRISPR guide target indicated in red. Sequence of CRISPR induced mutations below where red characters are guide target sequence and blue represents indels in both alleles. Gray box on exon 3 translation indicates CLE domain while blue characters indicate frame shifted amino acid sequence of *lsclv3-cr*. b-c, RT-qPCR of *GhCLV3* and *GhWUSb* transcripts in WT and *GhCLV3* RNAi Gerbera inflorescence meristems normalized to actin with mean and SEM (RQ 2^-ΔCq^) and biological replicates shown as individual dots. d, forest plot of RT-qPCR pairwise comparisons with 95% CI of mean difference (ΔCq_2_-ΔCq_1_); effect sizes reported as Hedge’s g in Supplementary File 8. Filled circle indicates p < 0.05 and unfilled indicates n.s (p > 0.05). e, petal and stigmata counts of WT vs *lsclv3-cr* florets. Statistics from Mann-Whitney U tests (p-values <0.0001). f, WT and *lsclv3-cr* capitulescence meristems of 4-week-old plants. g, representative images of WT and *lsclv3-cr* florets compared to aberrant terminal floret occurring in the center of all *lsclv3-cr* capitula. h, 5-week-old WT and *lsclv3-cr* plants. i, Stem widths of WT and *lsclv3-cr*. Statistics from Welch’s t-test (p-value < 0.0001). j, 38-day-old WT and ls*clv3-cr* plants with leaves removed to view primary axis. Roman numeral indicate apices resulting from aberrant splits in the primary meristem rather than axillary branching. Whole shoot, profile, and top-down images to scale across individuals. In b,c, n=3 biological replicates of pooled meristems from 3 or more individuals. In e, (n=individual plants with florets from 2 capitula counted each), n=10 (WT), 9 (*lsclv3-*cr). In i, (n=individual plants), n=27 (WT), 32 (*lsclv3-cr*). See complete statistical analyses and sample sizes in Supplementary File 8. Scale bars - 1mm.

**Supplementary Fig. 15:**
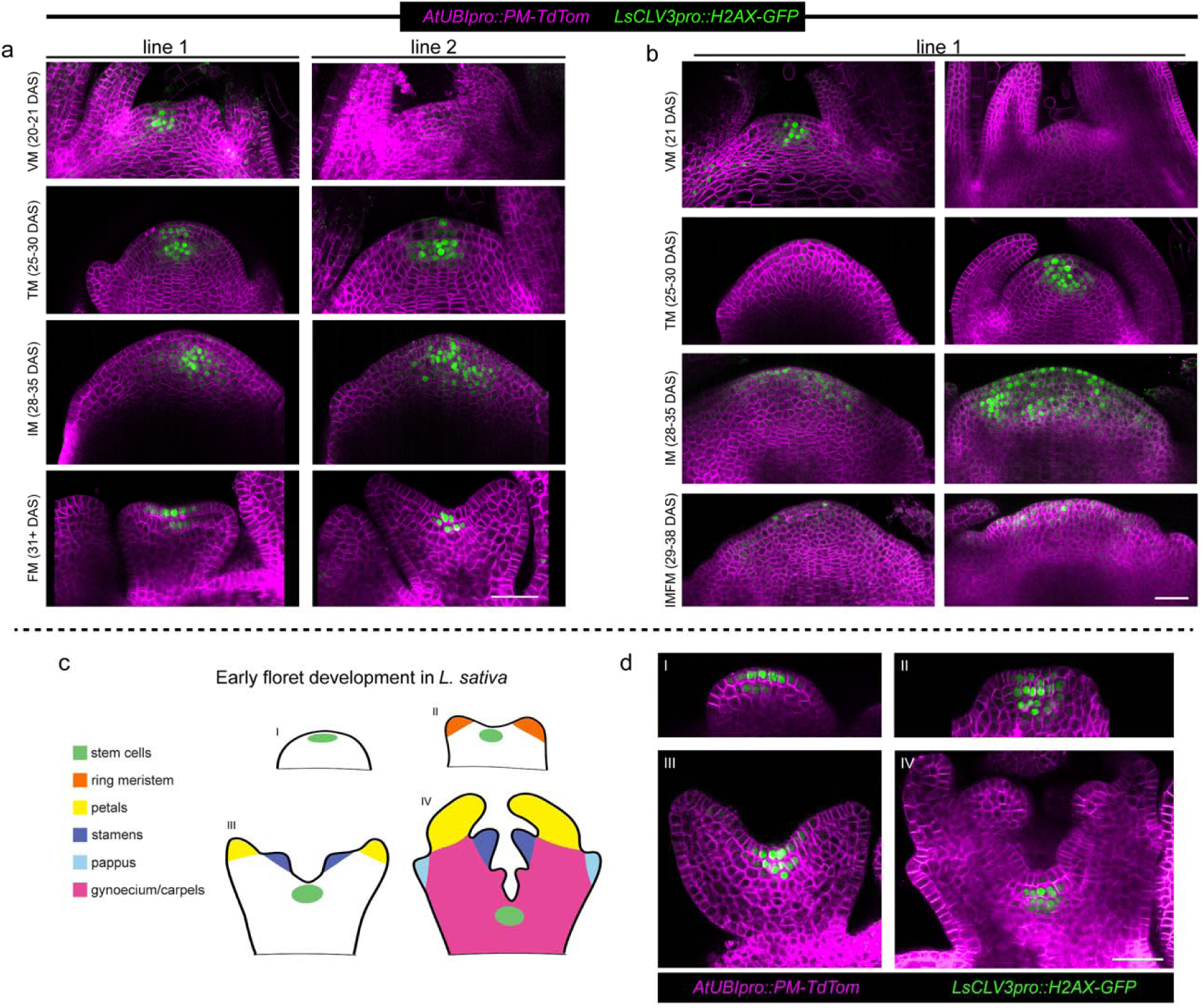
*LsCLV3pro* analysis across shoot, inflorescence and floral meristems. Confocal micrographs of *LsCLV3pro::H2AX-GFP* (green) and plasma membranes labeled by tdTomato with N-terminal fusion of the first 8aa of SlCPK1 (magenta). a, 20-21 day-old vegetative meristem, (VM), 25-30 day-old transition meristems (TM), 28-35 day-old inflorescence meristems (IM), and 31+ day-old floral meristems of two independently transformed lines represent between line variability. b, Two VMs, TMs, IMs, and 29-38 day-old inflorescence meristems with floral primordia (IMFM) from showing variability within a single transgenic line. c, A diagram of 4 stages of floral meristem development indicates the stem cell niche in green, the ring meristem (orange) which gives rise to petals (yellow) and stamen (purple) primordia, pappus primordia (blue), and developing gynoecium/carpels (magenta). d, micrographs of *LsCLV3pro* activity throughout floral development. Scale bars - 50µm.

**Supplementary Fig. 16:**
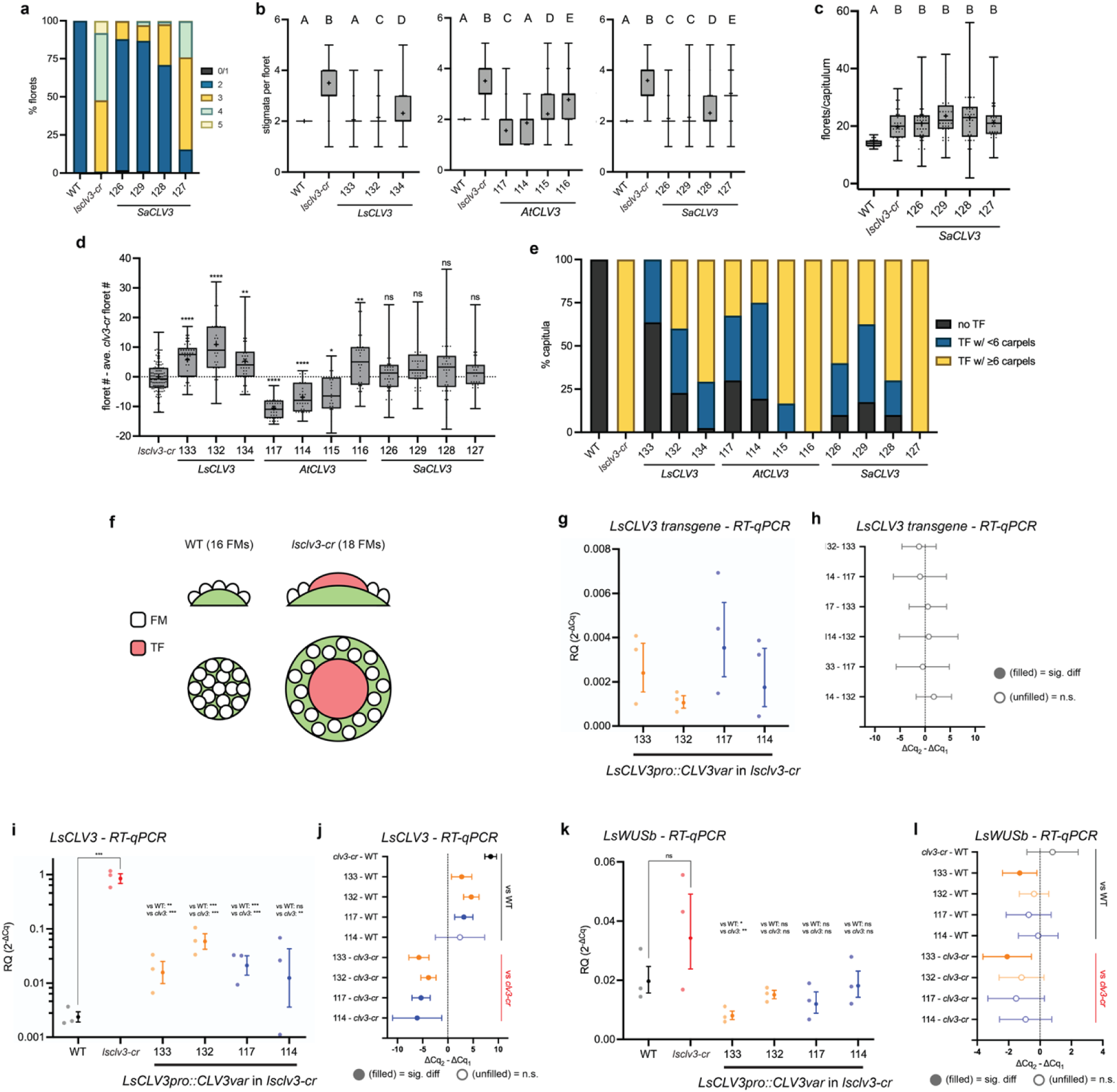
Functional analysis of *AsterCLV3* in lettuce. a, Complementation of increased stigmata number phenotype comparing *LsCLV3pro::SaCLV3* (lines 126, 129, 128, 127) expressed in *lsclv3-cr*. b, Box plots with statistical summaries from Kruskal-Wallis and Dunn’s multiple comparison tests from separate complementation experiments (related to Figure 5). c, Floret counts per capitulum (excluding terminal florets) comparing *LsCLV3pro::SaCLV3* to WT and *lsclv3-cr*. Statistical summaries from Kruskal-Wallis and Dunn’s multiple comparison tests where A and B groups are significantly different (adj. p-value <0.0001). d, Difference-in-differences plot comparing individual floret number per capitulum of *LsCLV3, AtCLV3,* and *SaCLV3* lines to the mean *lsclv3-cr* floret number in each respective experiment. Kruskal-Wallis with a Dunn’s multiple comparison test of *lsclv3-cr* across each of the 3 experiments indicate negligible batch effect (p > 0.3211). Statistical summaries from Welch’s ANOVA with Dunnett’s multiple comparison test comparing each sample to ls*clv3-cr* (* p < 0.05, ** p < 0.01, **** p < 0.0001). e, Bar charts summarizing percentage of capitula with no terminal floret, a terminal floret with < 6 carpels, and a terminal floret with ≥ 6 carpels. f, Illustration of WT and *lsclv3-cr* capitula with typical floral meristems (FM) in white and terminal floret meristems (TF) in red illustrating the impact of the terminal floret on the relationship between capitulum size and floret number. g-l, RT-qPCR of transgenic *LsCLV3* (g) transgenic and endogenous *LsCLV3* (i-j), and *LsWUSb* normalized to *LsUBI10* with mean and SEM (RQ 2^-ΔCq^) and biological replicates shown as individual dots. g, Welch’s ANOVA found no significant difference between transgene expression levels (p > 0.269). i, A generalized least squares model with custom contrasts comparing everything to WT and *lscvl3-cr* a indicates a significant difference in all comparisons except 117 vs WT (** p < 0.01, *** p < 0.001; p = 0.705). k, The same GLS model applied to *LsWUSb* transcript levels indicates no significant difference in any comparison except 133 vs WT and 133 vs *lsclv3-cr* (p = 0.012 and 0.002, respectively). h, j, l, forest plot of pairwise comparisons with 95% CI of mean difference (ΔCq_2_-ΔCq_1_) estimated from each test; effect sizes reported as Hedge’s g in Supplementary File 8. Filled circle indicates p < 0.05 and unfilled indicates n.s (p > 0.05). In b, (n=individual plants with 4 capitula counted each), from left to right: n=8 (WT), 10 (*lsclv3-cr*), 11 (133), 9 (132), 11 (132), 11 (134), 9 (WT), 10 (*lsclv3-cr*), 10 (117), 9 (114), 6 (115), 9 (116), 10 (WT), 9 (*lsclv3-cr*), 10 (126), 10 (129), 10 (128), 9 (127). In d, (n=individual difference from mean *lsclv3-cr*), n=29 (*lsclv3-cr*), 11 (133), 9 (132), 11 (134), 10 (117), 9 (114), 6 (115), 9 (116), 10 (126), 10 (129), 10 (128), 9 (127). In e, (n=individual plants with 4 capitula counted each), n=27 (WT), 29 (*lsclv3-cr*), 11 (133), 9 (132), 11 (132), 11 (134), 10 (117), 9 (114), 6 (115), 9 (116), 10 (126), 10 (129), 10 (128), 9 (127). In g-l, n=3 biological replicates of pooled developing florets from 8 or more capitula from 2 individuals. See complete statistical analyses and sample sizes in Supplementary File 8.

**Supplementary Fig. 17:**
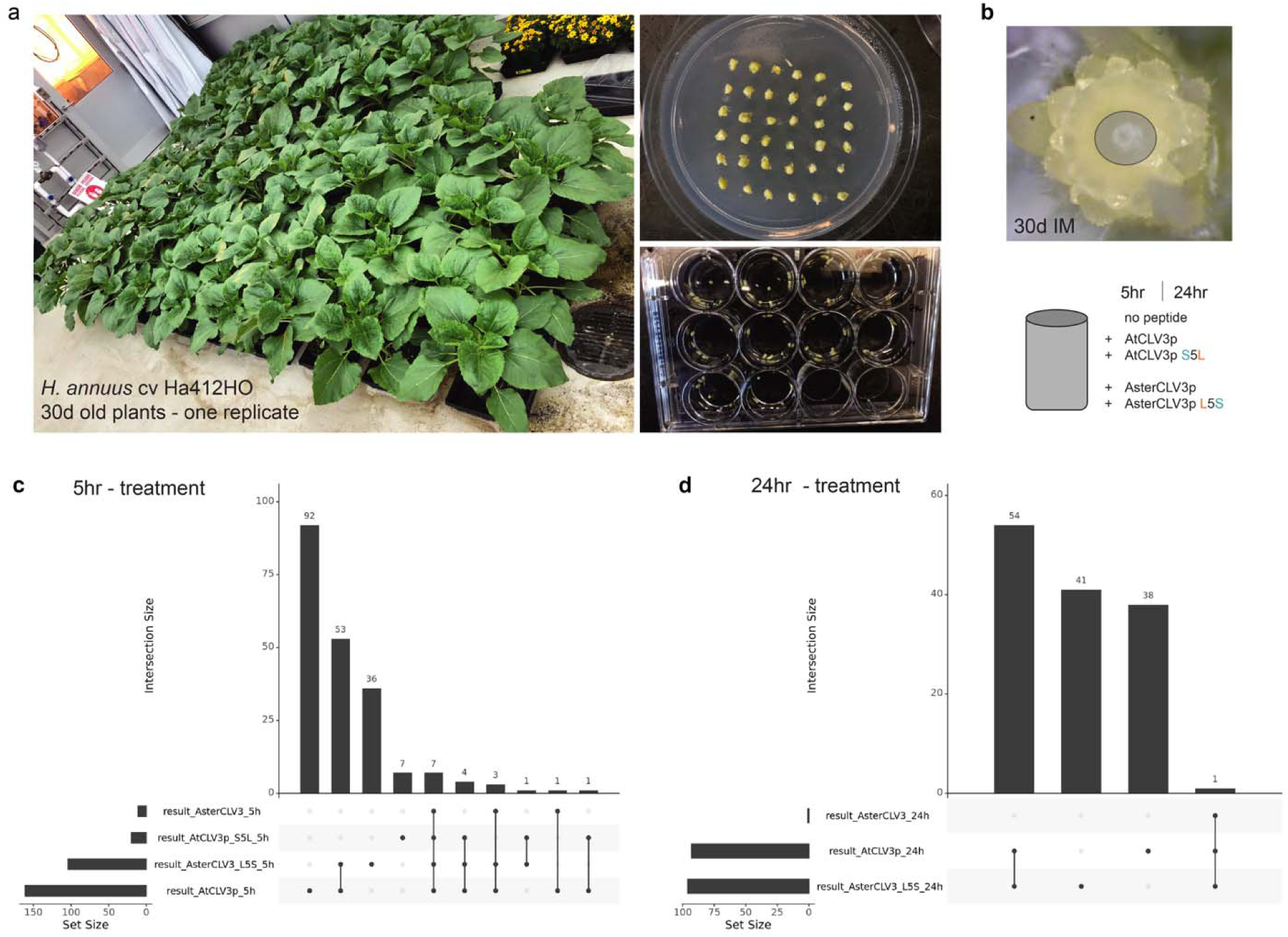
Transcriptomic analysis of CLEp-treated sunflower inflorescence meristems. a, 30-day old sunflowers (*H. annus* cv. Ha412HO). On right-top dissected shoot meristems at the inflorescence meristem (IM) expansion stage and bottom cored/isolated IMs in shoot culture media for CLEp treatments. b, op - representative stereoscope image of a 30d shoot meristem prior to tissue coring with circle indicating relative position of core sampling. Bottom - schematic of peptide treatment conditions before RNAseq. c-d, upset plots of differentially expressed (DE) genes across pairwise comparisons showing shared and unique DE genes in CLEp-treated sunflower IMs at 5hr (c) and 24hr (d) post-treatment.

**Supplementary Fig. 18:**
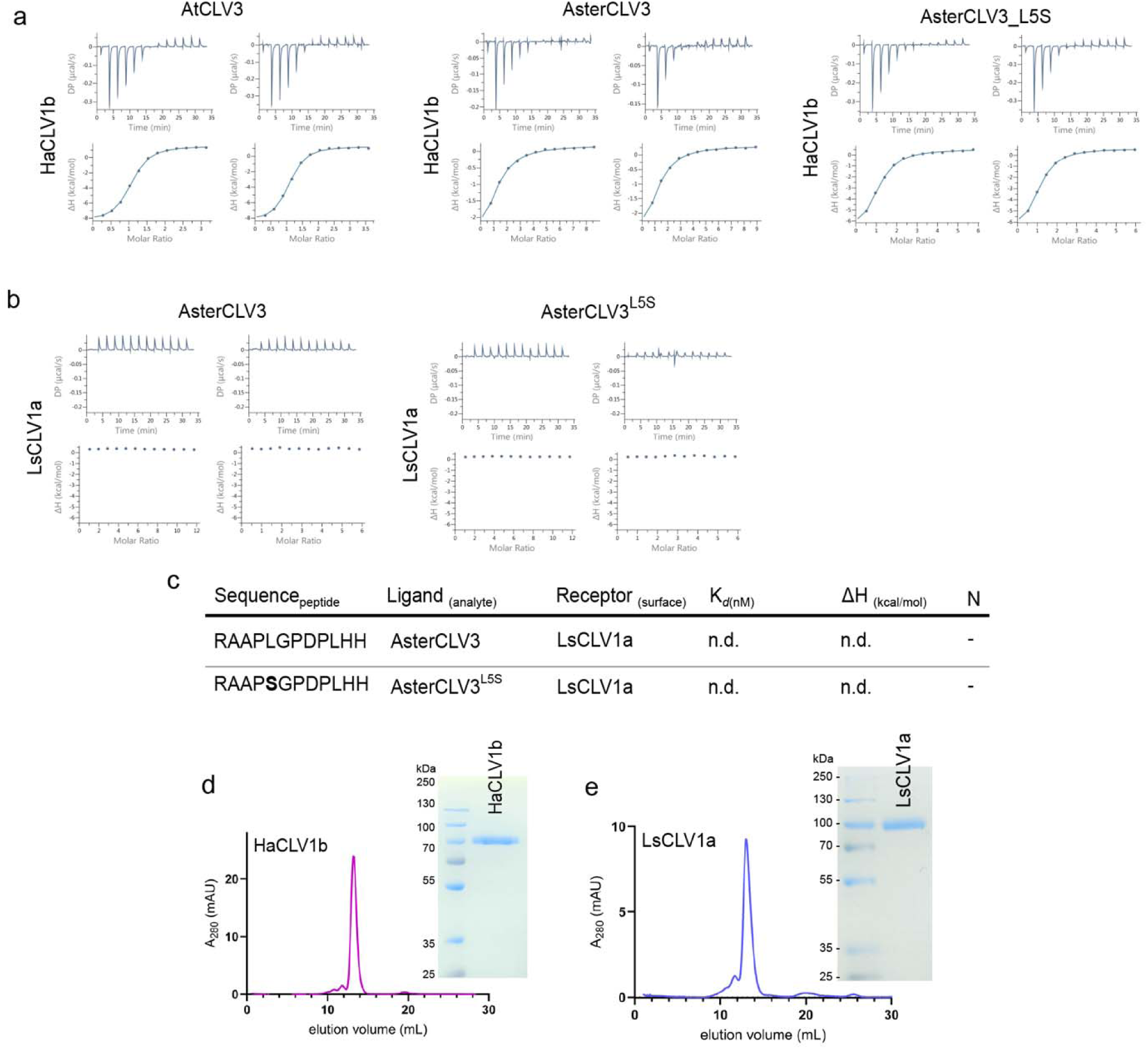
HaCLV1b and LsCLV1a peptide binding assays. a, Isothermal titration calorimetry (ITC) thermograms of HaCLV1b (sunflower) binding to different CLV/CLE peptides from independent experiments, corresponding to the analysis presented in Fig. 5k. b, ITC thermograms of LsCLV1a (lettuce) binding to different CLV/CLE peptides from independent experiments. c, ITC summary table of CLE/CLV peptides binding to the LsCLV1a ectodomain. K*_d_* (nM) represents binding affinity, ΔH (kcal/mol) indicates reaction enthalpy, and N denotes stoichiometry (n = 1 for a 1:1 interaction). Data are shown as non-detected binding (n.d.). d-e, Protein integrity and purity. SEC analysis and SDS-PAGE of HaCLV1b and LsCLV1a.

**Supplementary Fig. 19:**
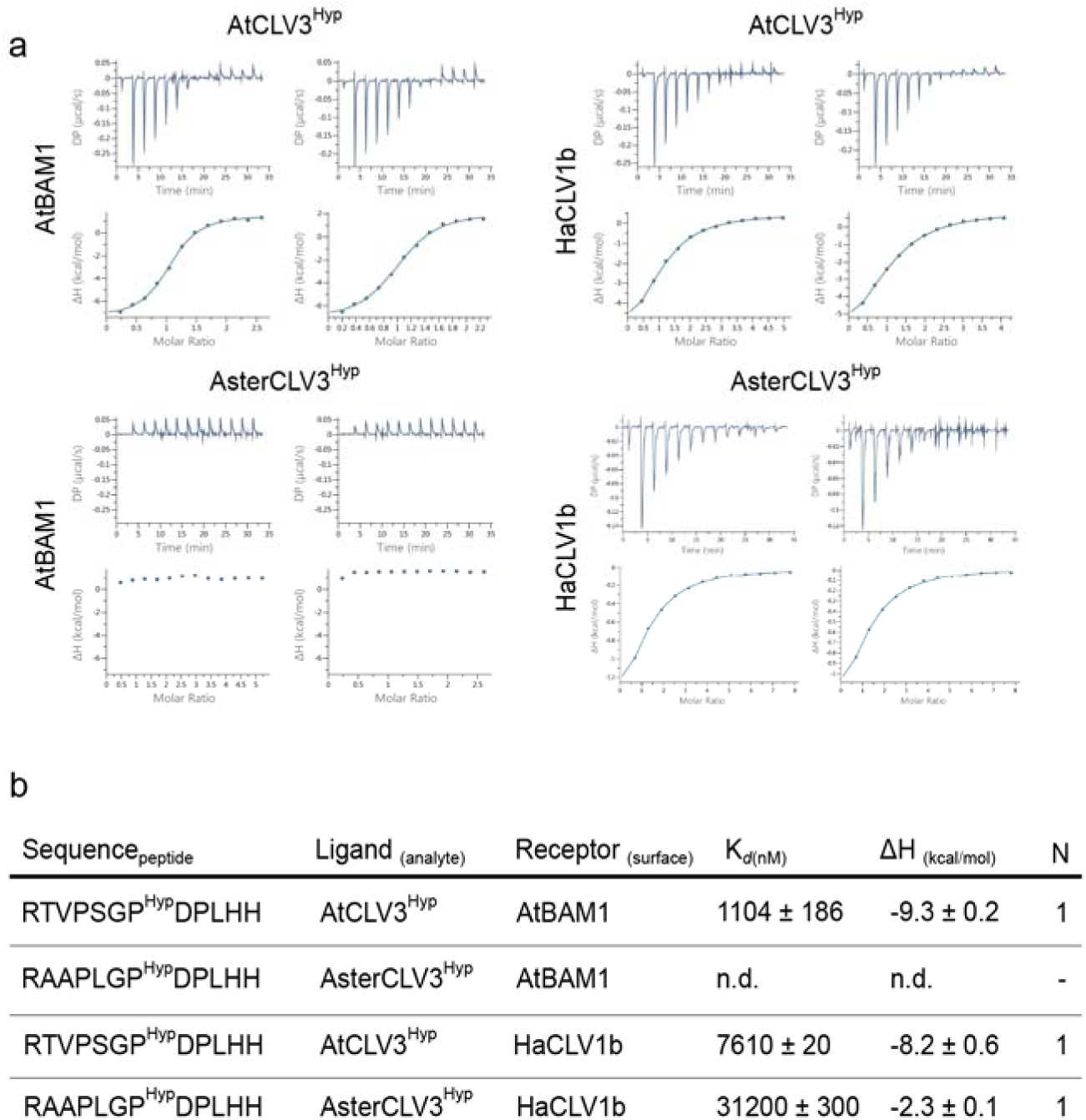
Binding of hydroxyprolinated CLV3 (CLV3Hyp) peptides to AtBAM1 and HaCLV1b. a, Isothermal titration calorimetry (ITC) analysis of hydroxyprolinated AtCLV3 and AsterCLV3 peptides binding to the AtBAM1 and HaCLV1b ectodomains. Representative thermograms from independent experiments are shown. b, Summary of ITC binding parameters. K*_d_* (nM), binding affinity; ΔH (kcal/mol), reaction enthalpy; and N, binding stoichiometry. Data are means ± SD from at least two independent experiments. n.d., no binding detected.

